# Synovial Fluid Spatial Microrheology is Shaped by Extracellular Matrix Identity and Mucin Integrity

**DOI:** 10.64898/2026.11.21.768317

**Authors:** Diego R. Jaramillo Pinto, Nathan C. Roehrig, Nicolas L. Mendoza, Matthew J. Paszek, Heidi L. Reesink, Lawrence J. Bonassar, George D. Degen, Roberto C. Andresen Eguiluz

## Abstract

The interfacial organization of synovial fluid (SF) contributes to cartilage protection and lubrication, yet how articular cartilage extracellular matrix (AC-ECM) identity influences the rheological response of supramolecularly assembled SF films remains to be elucidated. Here, depth-resolved multiple-particle tracking microrheology was used to examine diluted SF (dSF) near immobilized collagen type II (Col-II), collagen type I (Col-I), and fibronectin (FN). Untreated dSF exhibited increased relative apparent viscosity near Col-II and FN, whereas the response near Col-I was comparatively negligible. Treatment with StcE, a mucin-selective protease previously shown to cleave lubricin’s mucin domain in SF, resulted in more pronounced effects of the interface on the depth-dependent viscosity profiles across all three substrates. Analysis of the complex modulus and viscosity revealed significant effects with depth changes and StcE treatment. Quartz crystal microbalance measurements further revealed that the adsorbed amount of the fluids onto AC-ECM substrates did not directly correspond to the magnitude of microrheological modulation, with Col-I showing limited retention despite a pronounced near-interface response following StcE treatment. Together, these findings indicate that AC-ECM identity and StcE-sensitive SF constituents are associated with spatial variations in viscoelasticity that are not directly reflected by far-field tracer dynamics or interfacial adsorption alone.

## 1 Introduction

Synovial fluid (SF) is a viscoelastic, multicomponent biological fluid that contributes to the exceptional wear resistance and low friction of articulating cartilage surfaces.[1–3] Its mechanical and lubricating properties, that depend on confinement, normal loading, and shear rate,[2, 4–6] emerge from interactions among glycoproteins such as lubricin (LUB),[1, 7] glycosaminoglycans such as hyaluronic acid (HA),[4, 8–10] and phospholipids such as phosphatidylcholine (PC).[8, 10] At the articular cartilage extracellular matrix (AC-ECM) surface, SF constituents adsorb and assemble into hydrated interfacial films that contribute to boundary lubrication and protect the cartilage surface against wear.[11] The lubricating properties of these films are closely associated with molecular hydration and the ability to retain water under load.[12, 13] These mechanisms highlight the importance of interfacial molecular organization, which is not necessarily reflected in the properties of the bulk SF.[5] Consequently, bulk rheological characterization alone cannot fully describe the mechanical environment near the cartilage surface, where SF constituents interact with the AC-ECM surface. As an example of heterogeneous micro-rheological properties in SF-derived systems, HA–recombinant lubricin showed mechanically distinct microphases.[14] These findings highlight the value of spatially resolved tracer-scale measurements in complex fluids. Previous rheological work has also shown that the contribution of SF proteins must be distinguished from effects arising from artifacts at the rheometer interface, with HA dominating steady-shear viscosity for model SF composed of HA, albumin and γ-globulin (also present in native SF).[15] Together, these observations further motivate examining the response of multicomponent SF adjacent to molecularly defined solid interfaces rather than inferring its interfacial behavior from bulk rheology alone.

The molecular composition of the AC-ECM changes during injury, repair, and degeneration. Collagen type II (Col-II) is the most abundant collagenous component of healthy articular cartilage, whereas increased collagen type I (Col-I) expression and deposition has been associated with fibrocartilaginous repair and osteoarthritic changes.[16–18] Fibronectin (FN) is a distinct non-collagenous glycoprotein of the AC-ECM whose abundance, spatial distribution, and isoform composition is altered during osteoarthritic degeneration.[19, 20]

Recent studies have identified substantial differences in the assembly of SF-derived films on model AC-ECM substrates. Substantial retention of SF constituents was observed on Col-II and FN, whereas retention on Col-I was comparatively modest.[21, 22] Such behavior cannot be explained solely by the affinity of an individual boundary lubricant: recombinant equine lubricin (rEqLub), for example, was retained to comparable extents on Col-II and Col-I films.[21, 23] These observations are consistent with AC-ECM-dependent cooperative interactions among several SF constituents. However, characterizing a material adsorbed to an interface does not establish how its presence influences the mechanical response of the surrounding fluid, nor the spatial extent of such effects. Multiple-particle tracking microrheology (MPTM) is a technique that enables these variations to be probed. For example, it has been used to reveal increased apparent viscosity of HA near lubricin-intact cartilage, with an effect reported over distances extending approximately 50 µm from the surface.[24] Removal of native lubricin reduced this effect, whereas sequential incubation in SF restored the spatial profile.[24] Cartilage interactions have also been reported to increase the apparent viscosity of HA solutions relative to conventional surfaces.[5] Together, these studies establish that interactions with cartilage can modify measured lubricant rheology and spatial tracer dynamics. Nevertheless, they do not isolate the contribution of distinct AC-ECM molecular identities under otherwise comparable model-interface conditions. Understanding the contribution of SF macromolecular constituents to these spatial responses is critical to guide the design of intra-articular therapies aimed to recover SF function. LUB, a major boundary-lubricating glycoprotein in SF, contains a mucin-like domain that is heavily O-glycosylated and contributes to interfacial lubrication.[25, 26] The mucin-selective metalloprotease StcE has previously been shown to cleave lubricin’s mucin domain in SF, substantially impairing cartilage lubrication without producing a significant change in bulk viscosity under the conditions examined.[27] Although StcE can also cleave other mucin-like glycoproteins, its established activity toward lubricin provides a path to investigate the potential contribution of mucin-like molecules to SF mechanical behavior. Whether this perturbation influences the spatial microrheological response of SF near distinct AC-ECM interfaces remains unknown.

In this work, it was hypothesized that SF spatial microrheology would be altered by preferential adsorption onto molecularly distinct AC-ECM surface models. Furthermore, it was hypothesized that the response of StcE-treated SF would differ compared to untreated SF. To test this hypothesis, depth-resolved MPTM was used to investigate the spatial microrheological response of untreated and StcE-treated diluted SF (dSF) near immobilized Col-II, Col-I, and FN (schematized in Figure 1). Recombinant equine lubricin (rEqLub) was used as a simplified reference for the far-field tracer-scale response, while quartz crystal microbalance with dissipation (QCM-D) measurements were used to assess whether AC-ECM-dependent adsorbed mass could be related to differences in spatial microrheology profiles. Untreated dSF exhibited greater near-interface apparent-viscosity modulation adjacent to Col-II and FN than to Col-I. Following StcE treatment, the normalized near-interface responses became stronger and more similar across the three substrates, despite differences in their dSFs adsorbed mass densities. These findings reveal that AC-ECM identity and StcE-sensitive SF constituents regulate SF microrheology near AC-ECM model surfaces.

**Figure 1.**
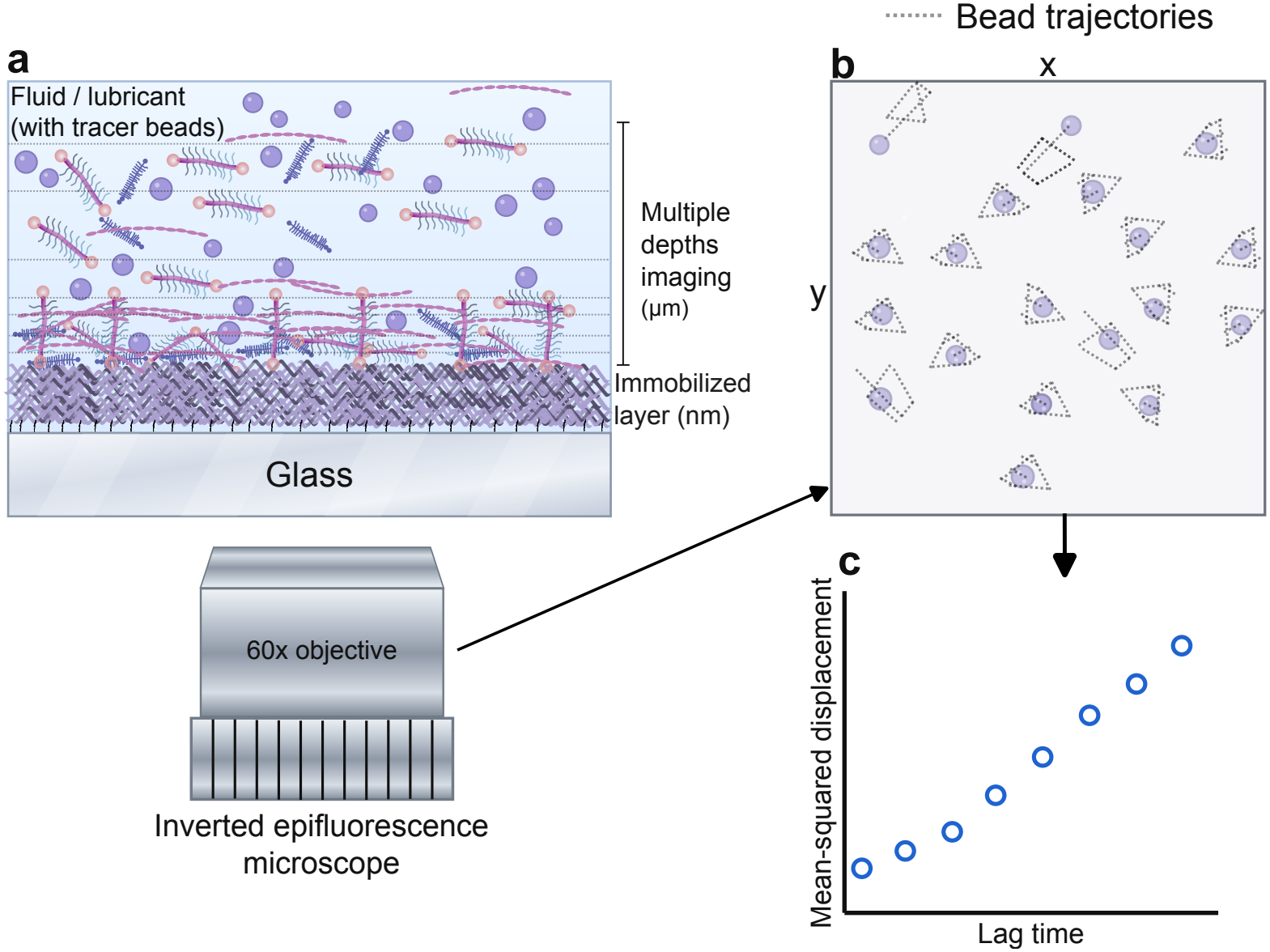
Depth-resolved microrheology at molecularly defined ECM interfaces. Schematic representation of depth-resolved multiple-particle tracking microrheology (MPTM). **a**, Fluorescent tracer particles were dispersed in the lubricant above an AC-ECM-functionalized glass substrate and imaged at defined distances from the interface using an inverted epifluorescence microscope. **b**, At each imaging depth, tracer-particle motion was recorded in the *x*-*y* plane and individual trajectories were reconstructed from sequential image frames. **c**, Bead trajectories were used to calculate lag-time-dependent MSDs and estimate depth-dependent microrheological properties.

## 2 Results and Discussion

### 2.1 Far-Field Microrheology Establishes Distinct Responses of dSF, StcE-dSF, and rEqLub

Far-field microrheology was first measured for each lubricant as a reference for the depth-dependent analysis. MPTM was performed at 100 µm above bare glass using 25% (v/v) SF in PBS (dSF), 250 µg/ mL rEqLub in PBS, and StcE-treated dSF (StcE-dSF). These measurements compared tracer-scale motion and frequency-dependent viscoelastic responses across the three lubricants without an immobilized AC-ECM surface model (Figure 2). These measurements should not be treated as an equivalent to bulk rheometry measurements.

**Figure 2.**
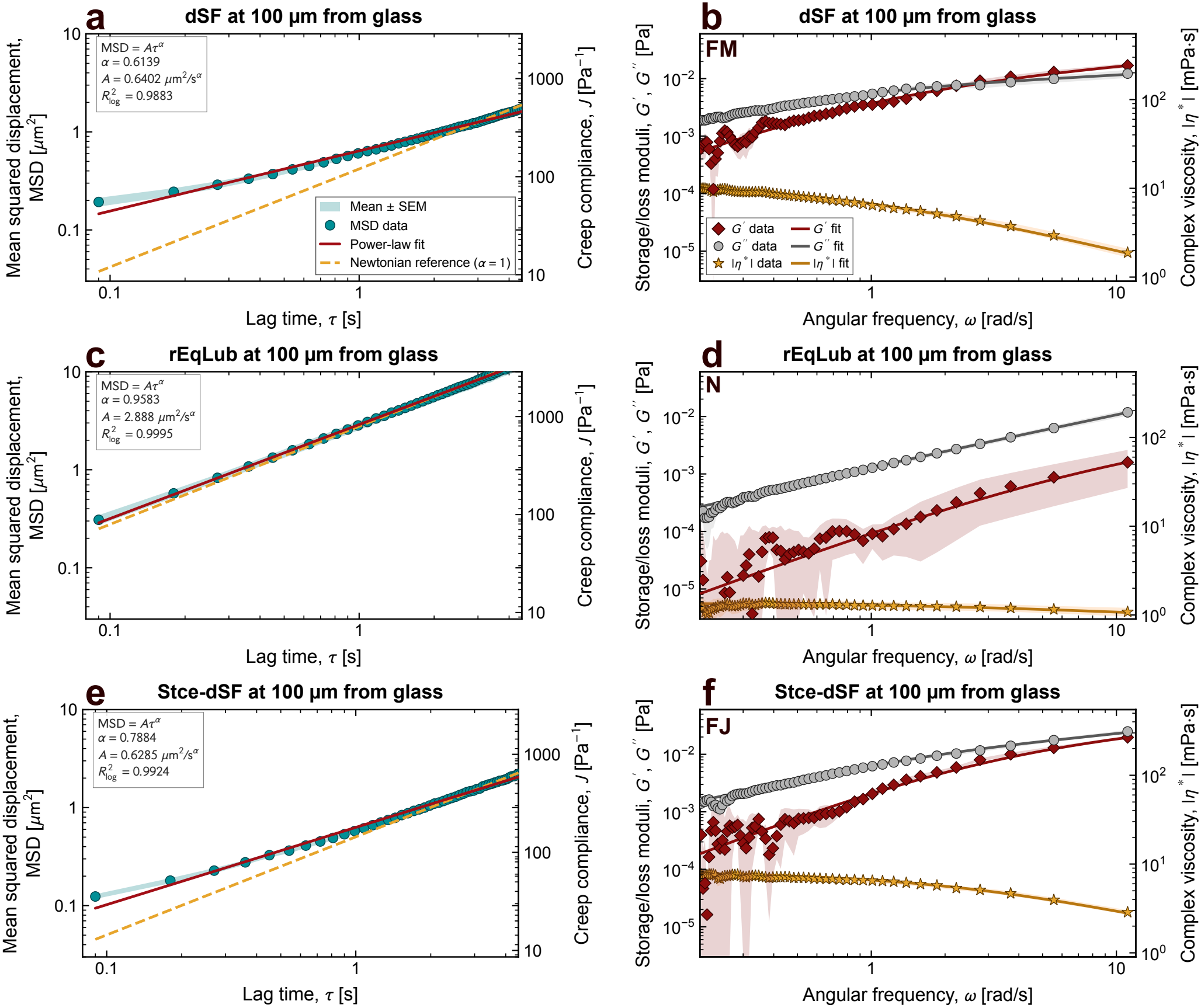
Far-field microrheology of synovial lubricants. **a**,**b**, Mean-squared displacement (MSD) and frequency-dependent storage modulus (*G*′), loss modulus (*G*″), and complex-viscosity magnitude (|η*|) of 25% (v/v) synovial fluid in PBS (dSF). **c**,**d**, Corresponding measurements for 250 µg /mL recombinant equine lubricin (rEqLub) in PBS. **e**,**f**, Corresponding measurements for StcE-treated dSF. All measurements were performed 100 µm from bare glass. In MSD panels, creep compliance (*J*) is shown on the secondary axis, with red solid lines indicating power-law fits (MSD = *A*τ^α^) and orange dashed lines indicating Brownian references (α = 1), anchored to the fitted MSD at the largest lag time included. In frequency-dependent panels, symbols represent experimental means, shaded regions indicate standard errors of the mean (SEM), obtained from at least 3 independent samples, and solid lines indicate the selected constitutive-model fits: fractional Maxwell liquid (FM) for dSF, Newtonian fluid (N) for rEqLub, and fractional Jeffreys-type (FJ) for StcE-dSF.

The mean-squared displacement (MSD) of tracer particles in dSF exhibited a subdiffusive dependence on lag time, with a fitted exponent of α_fit_ ≈0.61 (Figure 2a), consistent with previously reported values for full bovine SF.[1] The exponent below unity indicates restricted tracer motion over the fitted interval. The corresponding creep compliance, *J* (τ), displayed the same lag-time dependence as it is proportional to the MSD under the generalized Stokes–Einstein framework.[28] Here, the single power-law exponent, α_fit_, summarizes the MSD over a specified lag-time range, whereas the frequency-dependent analysis uses a local logarithmic exponent, α (τ), as described in the Experimental Section. Frequency-dependent measurements further distinguished the dissipative and elastic components of the dSF response (Figure 2b). At lower angular frequencies, the loss modulus exceeded the storage modulus (*G*″ > *G*′). With increasing frequency, *G*′ increased more strongly and approached *G*″; the selected fractional-Maxwell fit yielded a crossover near ω_c_ ≈ 2.82 rad/s within the analyzed interval. The magnitude of the complex viscosity, |η*|, decreased with angular frequency, indicating a frequency-dependent thinning. These observations are consistent with the viscoelastic properties of SF.[1, 29]

In contrast, tracer particles in rEqLub displayed substantially larger displacements and an approximately linear MSD, with α_fit_ ≈0.96 (Figure 2c). Consistently, the frequency-dependent response was predominantly dissipative, with *G*″ exceeding *G*′ throughout the measured interval and no experimentally resolved crossover (Figure 2d). The complex-viscosity magnitude showed little-to-no frequency dependence. Thus, purified rEqLub at the concentration examined behaved as a Newtonian fluid at the tracer scale, as reinforced by the optimally identified constitutive fit model (Newtonian). This far-field measurement provides a simplified compositional comparison with dSF. Studies of reconstructed HA–recombinant lubricin systems have identified heterogeneous microstructures, with mechanically distinct lubricin-rich aggregates dispersed within an HA-rich continuous phase.[14] However, the presence and mechanical properties of comparable aggregates were not established for the present rEqLub-in-PBS preparation.

StcE-dSF showed behavior intermediate between untreated dSF and rEqLub in its MSD scaling, with α_fit_≈ 0.79 (Figure 2e). Tracer motion therefore remained sub-diffusive after treatment but became closer to unconstrained Brownian scaling than in untreated dSF. The storage and loss moduli of StcE-dSF increased with frequency; *G*″ remained higher than *G*′ throughout the measured interval, and no modulus crossover was resolved (Figure 2f). The complex viscosity decreased with frequency, similar to untreated dSF. StcE treatment was therefore associated with an altered, but not eliminated, viscoelastic response of dSF. These changes are consistent with a contribution from StcE-sensitive SF constituents, including LUB, whose mucin domain has previously been shown to undergo StcE-mediated cleavage in SF.[27] Interestingly, previous studies reported substantial impairment of cartilage lubrication following StcE treatment without a corresponding significant change in bulk viscosity.[27] The present differences in tracer-scale dynamics therefore provide a complementary perspective on the mechanical consequences of StcE treatment to SF.

Constitutive-model analysis further distinguished the frequency-dependent responses of the two dSF preparations (SI Excel file). At 100 µm above bare glass, untreated dSF was best described by the fractional Maxwell liquid (FM) model, with an effective series-dashpot viscosity of η_*p*_ ≈12.2 mPa s and a springpot exponent of β≈ 0.30. However, StcE-dSF was best described by the fractional Jeffreys-type (FJ) model, with a lower η_*p*_ of approximately 7.75 mPa s and an additional parallel viscous contribution η_*s*_ ≈0.62 mPa s. No *G*″ = *G*′ crossover was measured within the fitted frequency range, consistent with the predominantly dissipative response of StcE-dSF. Thus, StcE treatment modified not only tracer mobility relative to untreated dSF but also the balance of viscous and elastic contributions across the frequency spectrum, even in the absence of immobilized AC-ECM proteins. Mechanistically, LUB has been shown to form complexes with HA,[4, 14, 30] which after StcE treatment could be compromised, resulting in a weakened SF macromolecular network. This can be seen as a decrease by |η*| and the loss of the *G*″ = *G*′ crossover point within the measured range.

Fractional constitutive models have similarly been used to describe the heterogeneous viscoelasticity of SF lubricating macromolecules. A Cross model, mathematically equivalent to the FJ representation used here, was used to characterize the continuous HA-rich phase of reconstructed HA–LUB mixtures, while identifying mechanically distinct LUB-rich aggregates.[14] Increasing HA concentration was associated with a decrease in the springpot exponent β from approximately 0.50 to 0.40, consistent with an increasingly elastic response of the polymer network.[14] It should be noted that the solvent viscosity was fixed at approximately 0.89 mPa s in their analysis, whereas the parallel viscosity contribution was fitted independently in the present study. Studies of a gel-forming mucin, MUC5AC, have similarly shown that changes in mucin-network organization can substantially alter tracer-scale dynamics.[31] In this context, the treatment-dependent constitutive response observed here is consistent with a reorganization of the macromolecular interactions governing dSF microrheology. Taken together, these far-field measurements establish three distinct reference states: markedly subdiffusive dSF, nearly Brownian rEqLub, and StcE-dSF with intermediate MSD scaling and persistent frequency-dependent viscoelasticity. The differences also make it necessary to distinguish absolute changes in tracer mobility from changes in the relative spatial contrast within each lubricant preparation when comparing untreated and StcE-treated dSF near AC-ECM surfaces (Col-II, Col-I, and FN).

### 2.2 dSF Near-interface Microrheology is modulated by AC-ECM Molecular Identity

To determine how AC-ECM molecular identity influences the microrheological response of adjacent dSFs, MPTM was performed at defined distances from immobilized Col-II, Col-I, and FN films. High-resolution N 1s XPS confirmed APTMS functionalization of the glass substrates (Figure S1), while liquid-environment AFM was used to characterize the morphology and apparent nanomechanics of immobilized FN films (Figure S2). Corresponding AFM characterization of immobilized Col-II and Col-I have been reported previously.[21]

The depth-dependent apparent-viscosity estimate was normalized to a condition-specific far-field reference defined by measurements at 50 and 100 µm, yielding the relative apparent viscosity, η_rel_ (Figure 3a). This scalar describes a relative change in resistance to tracer motion under a Stokes–Einstein-based approximation. As tracer motion is subdiffusive in dSF and the probes were measured near solid boundaries, the parameter should not be interpreted as an intrinsic continuum viscosity.

The normalized spatial profiles revealed distinct responses among the dSF/AC-ECM interfaces. Near Col-II, η_rel_ reached approximately twice its far-field value at the smallest measured distances, whereas near FN, η_rel_ exhibited a comparable increase. η_rel_ near Col-I remained much closer to its far-field value (Figure 3a). To quantify the spatial extent of these responses, the profiles were fitted using phenomenological sigmoid functions, from which the characteristic distance *d*_95_ was determined (see Methods). The fitted *d*_95_ values were approximately 9.3 and 5.3 µm for dSF on Col-II and FN, respectively, indicating that the near-interface enhancements diminished over different spatial ranges. In contrast, dSF near Col-I exhibited no pronounced increase in viscosity. The fitting parameters and model comparisons are provided in the SI (Figure S3). These trends are qualitatively consistent with the previously reported higher mass retention of dSF on Col-II and FN when compared to Col-I.[21, 22]

**Figure 3.**
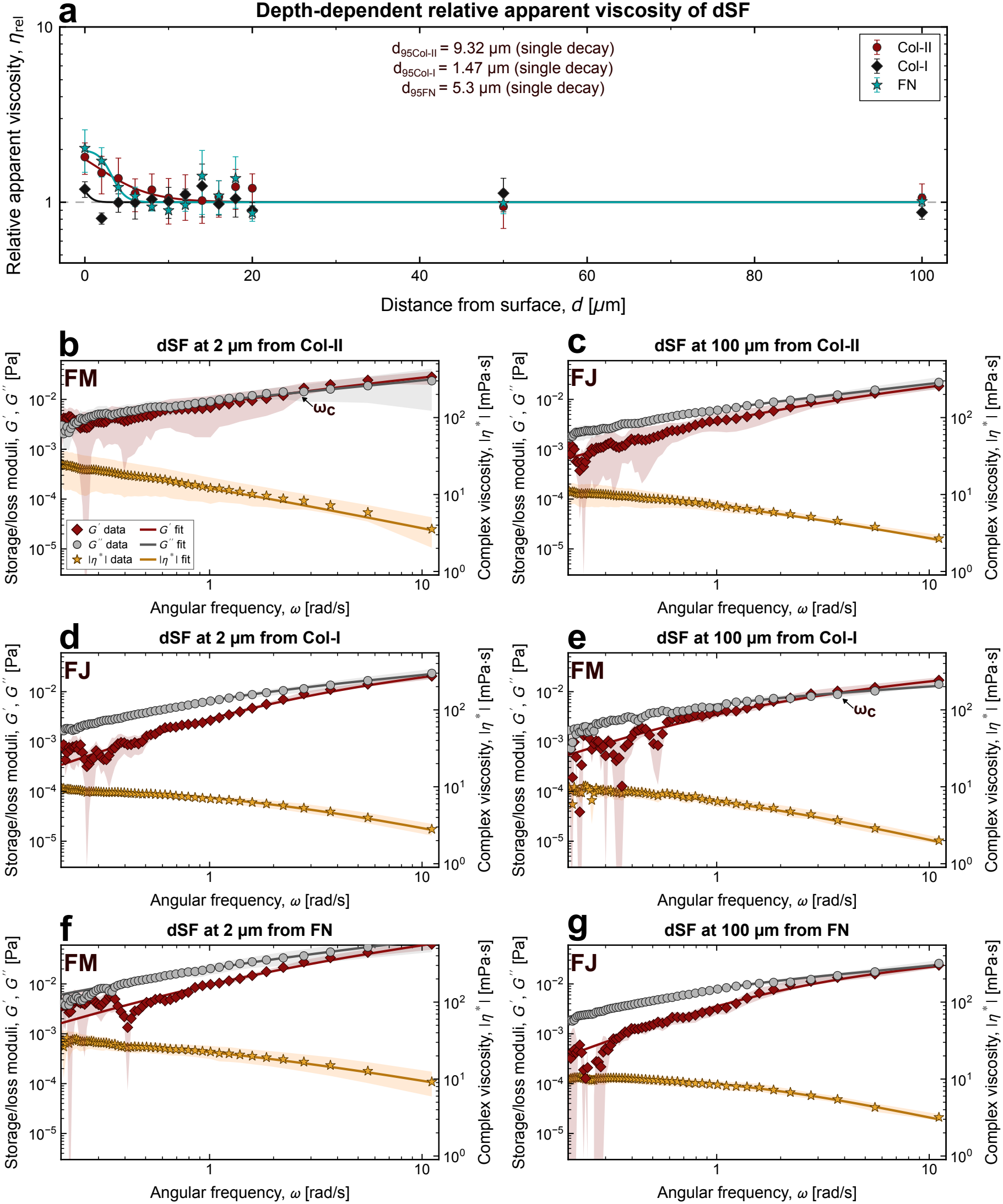
AC-ECM-dependent spatial microrheology of untreated diluted synovial fluid (dSF). **a**, Relative apparent viscosity, η_rel_, versus distance from immobilized Col-II, Col-I, and FN interfaces, calculated from tracer motion using a Stokes–Einstein-based approximation and normalized to the mean apparent viscosity at 50 and 100 µm for each AC-ECM/lubricant condition. Solid curves represent phenomenological single-Boltzmann spatial-decay fits, and fitted characteristic decay distances, *d*_95_, are also presented. **b–g**, Frequency-dependent storage modulus (*G*′), loss modulus (*G*″), and complex-viscosity magnitude (| η* |) at 2 and 100 µm from Col-II (**b**,**c**), Col-I (**d**,**e**), and FN (**f**,**g**). Symbols show experimental means, and solid lines represent selected constitutive-model fits. Moduli are plotted on the left axis and |η*| on the right. Error bars and shaded regions indicate standard errors of the mean (SEM) from at least three independent measurements.

Frequency-dependent microrheology was then evaluated at representative near-interface and far-field positions, 2 and 100 µm, respectively. Adjacent to Col-II, both dynamic moduli and complex viscosity were elevated at 2 µm relative to 100 µm, with the clearest differences below 1 rad/ s (Figure 3b,c). At 100 µm, the response was more dissipative over the majority of the frequency range, with the storage modulus approaching the loss modulus at higher frequencies. At both depths, |η*| decreased with increasing frequency. The spatial contrast near Col-II is therefore expressed not only in a scalar estimate of tracer resistance but also in the measured frequency-dependent response.

The near- and far-field spectra adjacent to Col-I were comparatively similar, although an apparent modulus crossover at 100 µm was not observed at 2 µm (Figure 3d,e). Some local differences, especially in the low-frequency storage-modulus estimates, remained visible, but the overall magnitudes and frequency dependencies were broadly comparable. This behavior was consistent with the relatively flat η_rel_ profile near Col-I and contrasted with the clearer increase measured near Col-II. These spectral differences suggest dissimilarities in molecular recruitment of dSF constituents. These findings are consistent with the broader literature establishing that cartilage-associated proteins can differ in the ways they retain and organize lubricating constituents.[6, 21– 23, 32, 33] Near FN, *G*′, *G*″, and |η*| at 2 µm generally exceeded their counterparts at 100 µm, and the storage contribution became more pronounced toward higher frequencies (Figure 3f,g). Thus, the untreated-dSF near-interface increase in apparent viscosity was not unique to Col-II, but its magnitude and frequency dependence varied among the three AC-ECM models. Unnormalized η (*d*) profiles for all interfaces, including bare-glass controls, are presented in SI (Figures S3–S6).

Constitutive-model fitting provided further evidence of AC-ECM-dependent differences in the frequency-dependent response (Table 1 and SI Excel file). At 2 µm, the fractional Maxwell liquid (FM) model was selected for dSF near Col-II and FN, whereas the fractional Jeffreys-type (FJ) model was preferred near Col-I. Interestingly, the model preferences differed at 100 µm, where FJ was selected for Col-II and FN, but FM for Col-I. Near Col-II, the selected FM fit at 2 µm predicted a crossover between *G*′ and *G*″ at approximately 2.43 rad /s, while no crossover was identified within the fitted interval at 100 µm. The opposite depth-dependence was observed near Col-I, where the selected far-field FM model predicted a crossover at 3.64 rad /s, which was absent from the near-interface FJ fit. Thus, the differences in tracer mobility among dSF/AC-ECM interfaces were accompanied by changes in the frequency-dependent viscoelastic response, rather than changes in modulus magnitude alone. This suggests differences in the partitioning of dSF components regulated by the underlying AC-ECM model protein films.

**Table 1.**
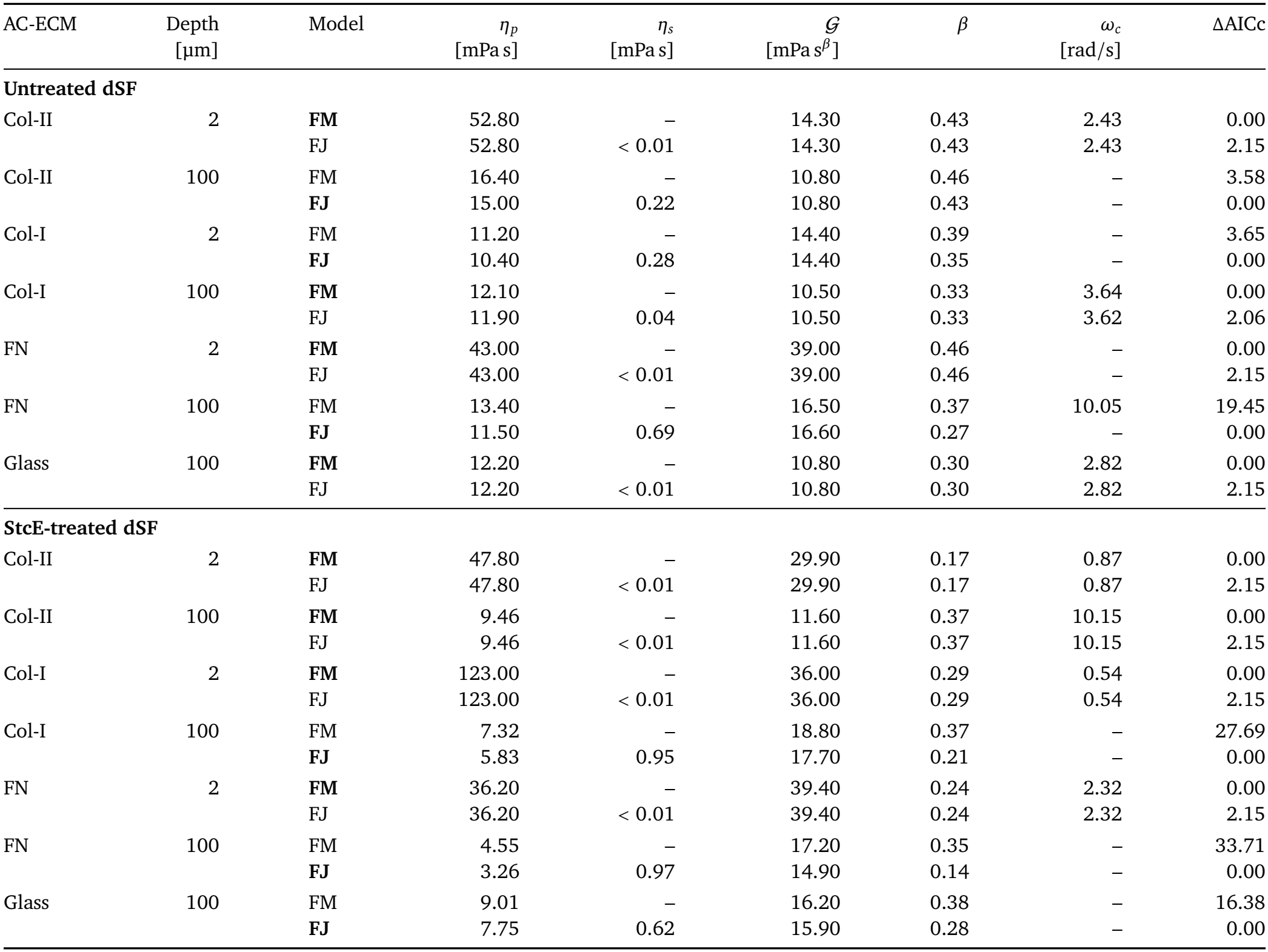
Comparison of the fractional Maxwell liquid (FM) and fractional Jeffreys-type (FJ) models used to fit dSF and StcE-dSF rheological properties, including fitted parameters and predicted *G*′ = *G*″ crossover frequencies (ω_*c*_). The model selected among all six candidates is bolded. ΔAICc is calculated relative to the lower-AICc model within each FM–FJ pair. A dash indicates no crossover within the measured range or a nonapplicable parameter. The parameters η_*p*_ and η_*s*_ denote the series and parallel dashpot viscosities, respectively, while *G* and β characterize the springpot. Expanded information on the parameters obtained from constitutive models can be found in SI (Excel file).

| AC-ECM | Depth | Model | *η<sub>p</sub>* | *η<sub>s</sub>* | 𝒢 | *β* | *ω<sub>c</sub>* | ΔAICc |
|  | [µm] |  | [mPa s] | [mPa s] | [mPa s<sup>*β*</sup>] |  | [rad/s] |  |
|---|---|---|---|---|---|---|---|---|
| **Untreated dSF** |  |  |  |  |  |  |  |  |
| Col-II | 2 | **FM** | 52.80 | – | 14.30 | 0.43 | 2.43 | 0.00 |
|  |  | FJ | 52.80 | < 0.01 | 14.30 | 0.43 | 2.43 | 2.15 |
| Col-II | 100 | FM | 16.40 | – | 10.80 | 0.46 | – | 3.58 |
|  |  | **FJ** | 15.00 | 0.22 | 10.80 | 0.43 | – | 0.00 |
| Col-I | 2 | FM | 11.20 | – | 14.40 | 0.39 | – | 3.65 |
|  |  | **FJ** | 10.40 | 0.28 | 14.40 | 0.35 | – | 0.00 |
| Col-I | 100 | **FM** | 12.10 | – | 10.50 | 0.33 | 3.64 | 0.00 |
|  |  | FJ | 11.90 | 0.04 | 10.50 | 0.33 | 3.62 | 2.06 |
| FN | 2 | **FM** | 43.00 | – | 39.00 | 0.46 | – | 0.00 |
|  |  | FJ | 43.00 | < 0.01 | 39.00 | 0.46 | – | 2.15 |
| FN | 100 | FM | 13.40 | – | 16.50 | 0.37 | 10.05 | 19.45 |
|  |  | **FJ** | 11.50 | 0.69 | 16.60 | 0.27 | – | 0.00 |
| Glass | 100 | **FM** | 12.20 | – | 10.80 | 0.30 | 2.82 | 0.00 |
|  |  | FJ | 12.20 | < 0.01 | 10.80 | 0.30 | 2.82 | 2.15 |
| **StcE-treated dSF** |  |  |  |  |  |  |  |  |
| Col-II | 2 | **FM** | 47.80 | – | 29.90 | 0.17 | 0.87 | 0.00 |
|  |  | FJ | 47.80 | < 0.01 | 29.90 | 0.17 | 0.87 | 2.15 |
| Col-II | 100 | **FM** | 9.46 | – | 11.60 | 0.37 | 10.15 | 0.00 |
|  |  | FJ | 9.46 | < 0.01 | 11.60 | 0.37 | 10.15 | 2.15 |
| Col-I | 2 | **FM** | 123.00 | – | 36.00 | 0.29 | 0.54 | 0.00 |
|  |  | FJ | 123.00 | < 0.01 | 36.00 | 0.29 | 0.54 | 2.15 |
| Col-I | 100 | FM | 7.32 | – | 18.80 | 0.37 | – | 27.69 |
|  |  | **FJ** | 5.83 | 0.95 | 17.70 | 0.21 | – | 0.00 |
| FN | 2 | **FM** | 36.20 | – | 39.40 | 0.24 | 2.32 | 0.00 |
|  |  | FJ | 36.20 | < 0.01 | 39.40 | 0.24 | 2.32 | 2.15 |
| FN | 100 | FM | 4.55 | – | 17.20 | 0.35 | – | 33.71 |
|  |  | **FJ** | 3.26 | 0.97 | 14.90 | 0.14 | – | 0.00 |
| Glass | 100 | FM | 9.01 | – | 16.20 | 0.38 | – | 16.38 |
|  |  | **FJ** | 7.75 | 0.62 | 15.90 | 0.28 | – | 0.00 |

A hydrodynamic contribution to the apparent-viscosity increase is expected because lateral tracer diffusion is hindered near a solid wall.[34, 35] For the nominal spherical tracer radius of 0.25 µm, the ideal planar, no-slip correction at a particle-center-to-wall separation of 2 µm corresponds to an estimated increase in lateral drag of approximately 7.5% (see Section S4). This correction is smaller than the approximately twofold normalized increase in apparent viscosity observed near Col-II and FN and cannot, by itself, explain the differences in tracer mobility measured at comparable nominal distances from the three substrates. The bare-glass controls (Figures S3– S6) show particularly large near-surface changes in tracer mobility, further highlighting the sensitivity of the measurement to the interface. Overall, these experiments revealed AC-ECM-dependent spatial differences in tracer mobility and the associated apparent viscoelastic spectra of dSF. Col-II and FN exhibited clearer near-interface effects than Col-I. Depth-dependent rEqLub measurements (Figure S5) showed pronounced near-interface reductions in tracer mobility, comparable to those observed in tracer-only PBS controls (Figure S6). These profiles indicate that rEqLub by itself does not assemble into thick enough layers to alter the surrounding fluid viscoelastic properties at the working resolution.

### 2.3 StcE Treatment Modifies Spatial Microrheology of dSF near AC-ECM Surfaces

The far-field measurements established that StcE treatment altered the diffusive and frequency-dependent response of dSF even without immobilized AC-ECM proteins (Figure 2). Given the previously demonstrated activity of StcE toward lubricin’s mucin domain in SF,[27] the same depth-resolved analysis was performed near Col-II, Col-I, and FN to examine how this mucin-directed perturbation influences the spatial response of the fluid (Figure 4). Relative to untreated dSF, StcE-dSF exhibited substantially greater normalized near-interface increases in apparent viscosity across all three substrates. At 2 µm from all three interfaces, η_rel_ reached approximately 3.5– 5.7 times the condition-specific far-field reference (Figure 4a). The profiles decreased over the first several micrometers, with fitted characteristic distances (*d*_95_; see Methods) of approximately 17.1, 9.9, and 8.0 µm for StcE-dSF on Col-II, Col-I, and FN, respectively. These distances were greater than those obtained for untreated dSF near Col-II (9.3 µm) and FN (5.3 µm), whereas on Col-I changed from a negligible response to a pronounced near-interface increase extending over approximately 10 µm. While the StcE-dSF profiles near Col-I and FN were adequately described by single-sigmoid functions, on Col-II the profile exhibited a weaker secondary shoulder extending farther from the interface, being better captured by a double-sigmoid model (Figure S4). This secondary contribution accounted for the larger fitted *d*_95_ near Col-II. Non-normalized values of η (*d*) and fitting-model comparions are presented in SI (Figures S3–S6). At 2 µm from Col-II, StcE-dSF displayed notable storage and loss contributions, with *G*′ increasing and exceeding *G*″ towards higher frequencies (Figure 4b). At 100 µm, the spectra were generally more dissipative, with smaller moduli, especially at lower frequencies (Figure 4c).

**Figure 4.**
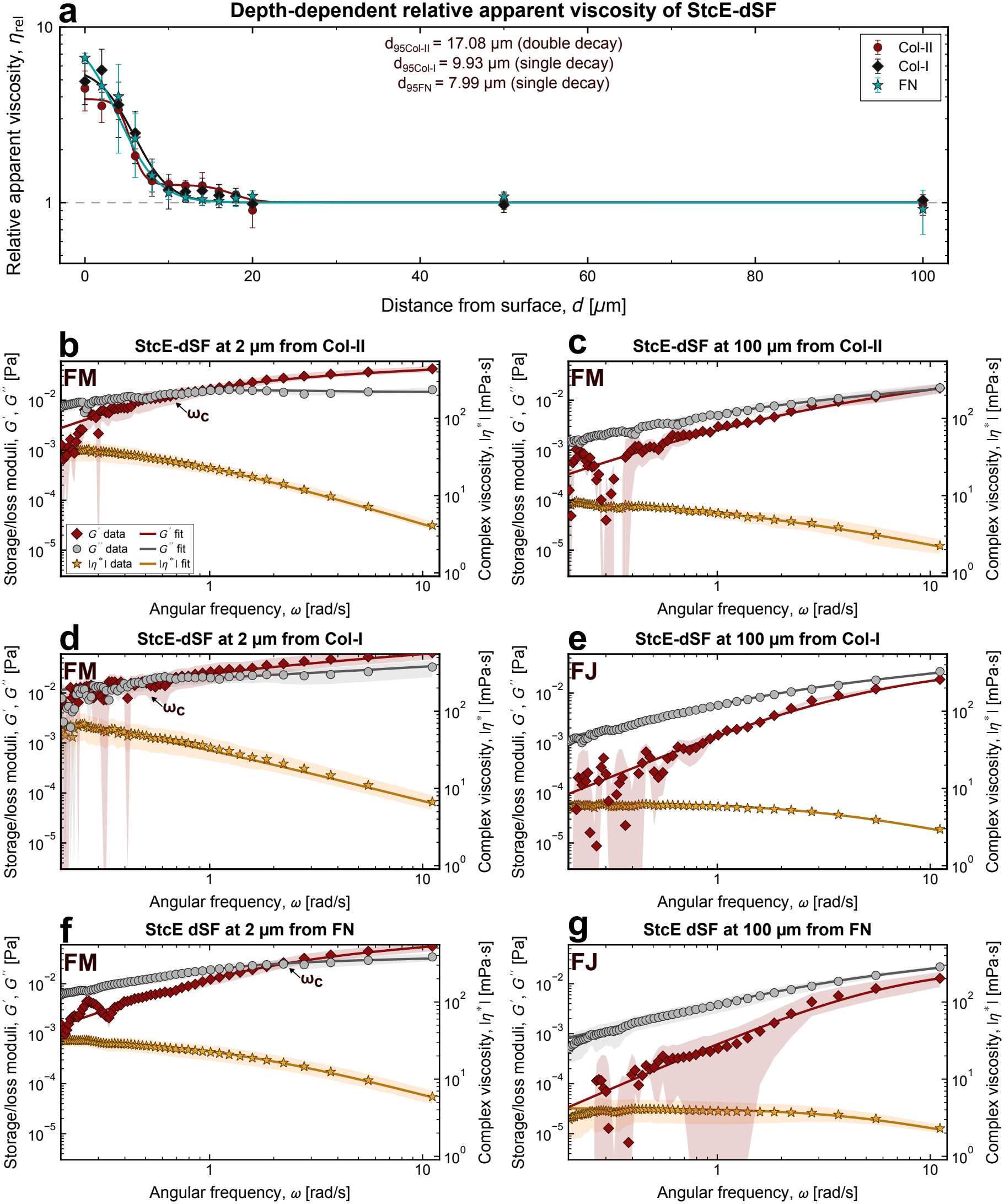
AC-ECM-dependent spatial microrheology of StcE-treated diluted synovial fluid (StcE-dSF). **a**, Relative apparent viscosity, η_rel_, versus distance from immobilized Col-II, Col-I, and FN interfaces, normalized to the mean apparent viscosity at 50 and 100 µm for each condition. Solid curves represent phenomenological single- or double-Boltzmann spatial-decay fits, and fitted characteristic decay distances, *d*_95_, are also presented. **b–g**, Frequency-dependent storage modulus (*G*′), loss modulus (*G*″), and complex-viscosity magnitude (|η*|) at 2 and 100 µm from Col-II (**b**,**c**), Col-I (**d**,**e**), and FN (**f**,**g**). Symbols show experimental means, and solid lines represent selected constitutive-model fits. Moduli are plotted on the left axis and |η*| on the right. Error bars and shaded regions indicate standard errors of the mean (SEM) from at least three independent measurements.

A prominent treatment-associated change emerged near Col-I. For dSF, the near-interface and far-field spectra were similar. After StcE treatment, the spectrum at 2 µm showed a stronger storage contribution, with *G*′ becoming comparable to or greater than *G*″ over part of the examined frequency range (Figure 4d). At 100 µm, *G*″ remained dominant (Figure 4e). Thus, a pronounced depth-dependent response appeared after StcE treatment even near Col-I, which had exhibited no normalized spatial modulation in untreated dSF (Figure 3).

A related near-to-far change was observed near FN. At 2 µm, *G*′ increased strongly with frequency, with a *G*″ = *G*′ crossover point ω_*c*_ = 2.32 rad/ s (Figure 4f). The 100-µm spectrum was more loss-dominated, particularly at lower frequencies (Figure 4g). The far-field *G*′ estimates in this condition were relatively scattered, however, and the low-frequency differences should not be over-interpreted. Across all three AC-ECM models, |η*| decreased with increasing frequency. StcE-dSF therefore retained frequency-dependent responses and AC-ECM surface specific viscoelastic modulation, even as the normalized scalar spatial profiles became more similar.

These spatial differences should be considered along-side the established functional effects of StcE: degradation of Lub’s mucin domain. Intact Lub has been reported to form complexes with HA, providing thick, viscous Lub and wear-protecting films.[24, 36] Compromised Lub integrity impairs cartilage lubrication substantially, without a corresponding significant change in bulk steady-shear viscosity.[27] Our findings elucidate that compromising SF mucins, in particular Lub, does not impair the supramolecular assembly of a SF-derived film. Furthermore, for the Col-I condition, it appears to favor the assembly of a film. Yet, the viscoelastic properties of these films differ from those derived from untreated SF.

The constitutive-model analysis revealed differences in the viscoelastic response of untreated dSF depending on both AC-ECM identity and distance from the interface (Table 1). At 100 µm, the FJ model was preferred for dSF exposed to Col-II and FN, whereas the FM model was selected for Col-I. Despite sharing the same preferred model, the Col-II and FN conditions exhibited different fitted parameters. In particular, the effective parallel viscosity, η_*s*_, was approximately 0.22 and 0.69 mPa s for Col-II and FN, respectively, while the springpot exponent β differed from 0.43 to 0.27. At 2 µm, FM was preferred for both Col-II and FN, although their responses remained distinct. Near Col-II, the fitted series-dashpot viscosity, η_*p*_, was approximately 52.8 mPa s, with a predicted *G*′ = *G*″ crossover at 2.43 rad /s, whereas near FN, η_*p*_ was approximately 43.0 mPa s, without a crossover within the experimental frequency range. In contrast, the selected FJ response near Col-I exhibited a comparatively small series-dashpot viscosity, consistent with the absent spatial dependence observed for untreated dSF on this substrate.

These differences are particularly interesting considering the adsorption and retention of SF constituents by the different AC-ECM interfaces. Although measurements at 100 µm were performed beyond the region of pronounced spatial viscosity enhancement, the fluid had nevertheless been exposed to the corresponding immobilized AC-ECM model. We hypothesize that selective adsorption and partitioning of SF constituents at these interfaces may influence the composition and organization of the remaining fluid, thereby contributing to the differences in its far-field rheological response. Our previous work demonstrated that Col-II effectively scaffolds SF-derived films, as opposed to Col-I.[21] Similarly, FN has been shown to support SF-derived film formation and to inter-act cooperatively with lubricin.[6, 22] Importantly, Col-II and FN may preferentially recruit different populations or relative proportions of SF constituents, even when both exhibit substantial adsorption. Such differences could alter the balance between freely dispersed macro-molecules and molecular assemblies, resulting in distinct constitutive parameters despite similar preferred model architectures. Closer to the interfaces, the additional organization of retained SF constituents could further account for the differences between the near-interface and far-field responses.

Following StcE treatment, the constitutive responses were reorganized in a distinctly AC-ECM-dependent manner. At 100 µm, the preferred model changed from FJ to FM near Col-II, with η_*p*_ decreasing from 15.0 to 9.46 mPa s and a predicted crossover emerging at 10.15 rad /s. Conversely, the model near Col-I changed from FM to FJ, with an effective parallel viscosity of approximately 0.95 mPa s. Near FN, FJ remained the preferred model, although η_*p*_ decreased from 11.5 to 3.26 mPa s, while η_*s*_ increased from 0.69 to 0.97 mPa s. These changes were accompanied by decreases in the springpot exponent for all three AC-ECM conditions, indicating a treatment-dependent modification of the fractional viscoelastic response. Interestingly, the QCM-D measurements showed that StcE-dSF retention remained substantial on Col-II, whereas retention on FN was slightly reduced relative to untreated dSF (Figure S7). Thus, the rheological changes did not simply follow the total amount of adsorbed material. Instead, mucin-domain cleavage may alter the interactions, relative affinities, and partitioning of different SF constituents at each interface, producing distinct changes in the remaining fluid. It is important to keep in mind that QCM-D and MPTM probe different spatial scales of the viscoelastic properties of the dSF/AC-ECM interfaces.

At 2 µm, StcE-dSF exhibited a more consistent constitutive response across the AC-ECM interfaces, with FM selected for Col-II, Col-I, and FN. For each condition, the additional parallel viscosity in the alternative FJ representation approached zero, indicating that the FM model adequately captured the measured spectra (Table 1). Nevertheless, the fitted parameters and crossover frequencies remained dependent on AC-ECM identity. The springpot exponent decreased from 0.43 to 0.17 near Col-II, from 0.35 to 0.29 near Col-I, and from 0.46 to 0.24 near FN. The predicted crossover near Col-II shifted from 2.43 to 0.87 rad/s, while crossovers at 0.54 and 2.32 rad/s were obtained for treated dSF near Col-I and FN, respectively. Notably, the fitted η_*p*_ near Col-I increased from approximately 10.4 to 123 mPa s, along with a pronounced near-interface enhancement that was largely absent for untreated dSF. Thus, although StcE treatment produced more similar normalized spatial profiles and a common preferred constitutive architecture at 2 µm, the mechanical responses retained clear AC-ECM-dependent characteristics.

Complementary QCM-D measurements further revealed AC-ECM-dependent adsorption and retention of StcE-dSF (Figure S7). Following rinsing, Col-II exhibited the greatest retained mass density, followed by FN and Col-I, with significant differences among all three substrates (Holm-adjusted *p* < 0.05). Comparison with previously reported untreated-dSF measurements showed no statistically significant treatment-dependent differences in apparent mass density on individual substrates.[21, 22] Interestingly, despite its limited retention on Col-I, StcE-dSF exhibited a pronounced near-interface viscoelastic response, including a substantial increase in the fitted η_*p*_ and enhanced storage and loss moduli relative to the far field.

Related studies of mucin-based systems have similarly revealed distinct relationships among interfacial adsorption, viscoelasticity, and lubrication. Changes in mucin molecular configuration can substantially alter lubricity with comparatively small changes in adsorption, while mucin networks with similar viscoelastic properties can exhibit different wear-protection behavior.[37, 38]

Together, these findings suggest that the microrheological environment near cartilage molecular models is influenced by two interconnected processes: the partitioning of SF constituents between the surface and the surrounding fluid, and their local organization in the immediate interfacial region. The first may contribute to the differences detected at far-field positions, whereas the second becomes particularly important at distances of a few micrometres from the interface. StcE-mediated perturbation modifies both contributions, changing the viscoelastic response of the fluid while also influencing how its constituents interact with different AC-ECM proteins. This interpretation is consistent with the heterogeneous rheology of reconstructed HA–Lub systems, in which the mechanical response depends on the relative organization of the constituent macromolecules.[14]

To further quantitatively compare the local viscoelastic responses, complement the constitutive-model fitting shown above, and give the findings a broader context, *G*″, *G*′, |η* |, and the loss tangent (tan δ = *G*″ *G*′) were evaluated and compared at ω = 1 rad/s, or equivalently *f*≈ 0.16 Hz, for each independent measurement (Figure 5). This frequency corresponds to the same nominal τ = 1 s timescale used for the scalar apparent-viscosity reference, although the two quantities are calculated differently. This low-frequency reference is also consistent with previous oscillatory model-SF rheology and falls within the frequency ranges examined in slow cyclic joint-flexion and contact-migration experiments.[15, 39, 40] This value has been used in biomechanical studies of gait.[41]

**Figure 5.**
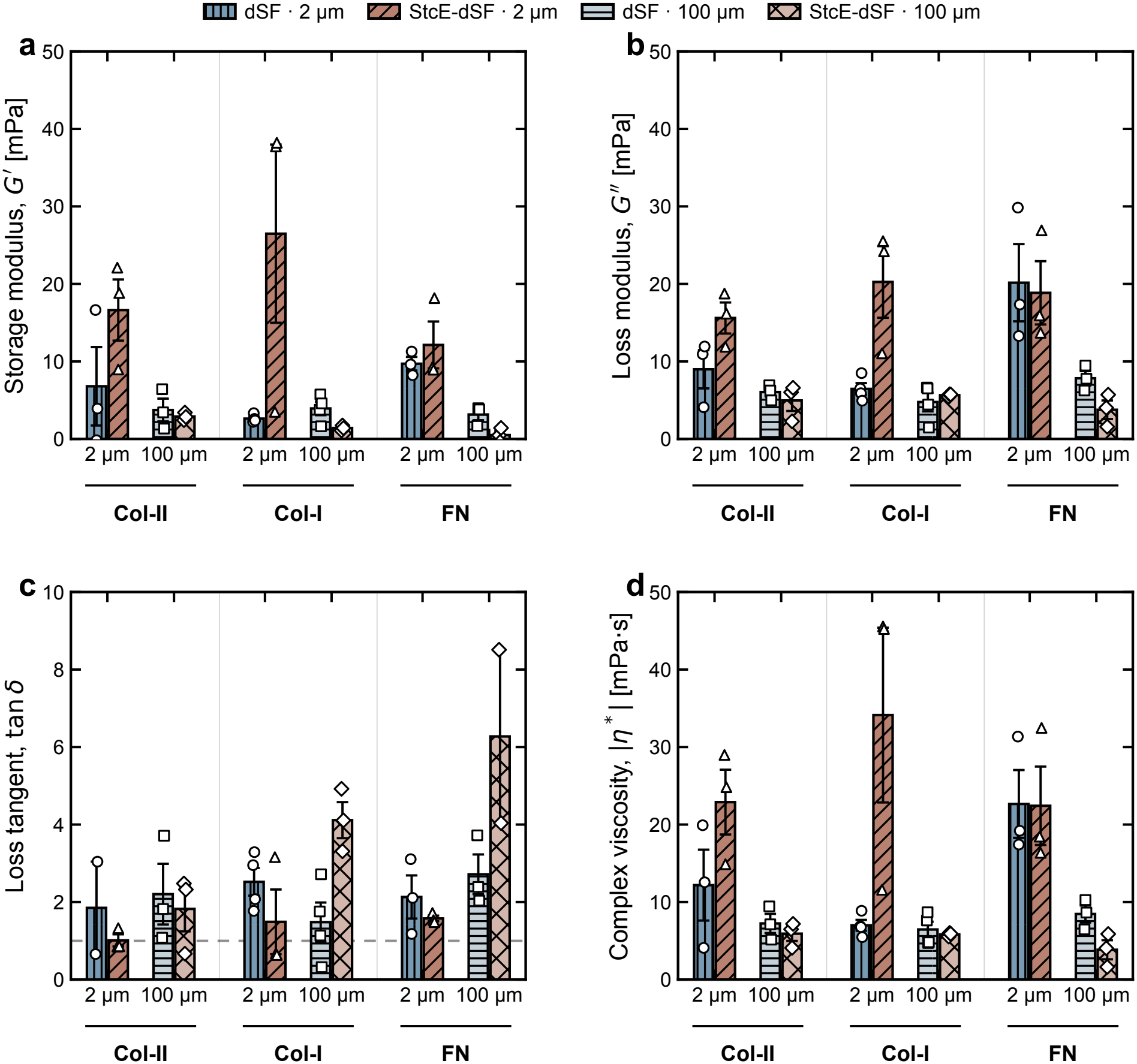
Effects of ECM identity and StcE treatment on spatial microrheology. **a–d**, Storage modulus (*G*′, **a**), loss modulus (*G*″, **b**), loss tangent (tan δ = *G*″/*G*′, **c**), and complex-viscosity magnitude (|η* |, **d**) at ω = 1 rad/s for dSF and StcE-dSF at 2 and 100 µm from Col-II, Col-I, and FN surfaces. In **c**, the dashed line indicates tan δ = 1. Bars show mean ± SEM; symbols represent independent measurements (*n* ≥ 3).

Across the AC-ECM interfaces, StcE-dSF generally exhibited larger near-interface moduli and complex viscosity than dSF, particularly near Col-I, whereas the corresponding values at 100 µm were comparatively similar between treatments. The loss tangent further distinguished the relative elastic and viscous contributions, with tan δ > 1 indicating a predominantly dissipative response, and tan δ < 1 indicating a predominantly elastic response. At 2 µm, StcE-dSF generally exhibited lower tan δ than dSF, approaching unity near Col-II and indicating a greater relative elastic contribution. In contrast, the far-field response remained predominantly dissipative, with particularly elevated tan δ for StcE-dSF near Col-I and FN. These differences highlight changes in the balance between elastic and viscous contributions that are not fully captured by the moduli or complex-viscosity magnitude alone. A three-factor split-plot ANOVA, accounting for paired measurements at different depths (2 and 100 µm), identified significant depth effects for *G*′, *G*″, and |η* | (*p* < 0.001, Table S1). Significant fluid treatment× depth interactions were also observed for *G*′ (*p* = 0.0046), *G*″ (*p* = 0.0134), and |η*| (*p* = 0.0056), indicating that StcE treatment modifies the spatial dependence of the viscoelastic response. These interactions remained significant under heteroscedasticity-robust sensitivity analysis. However, the three-way fluid treatment × AC-ECM × depth interactions were not significant, and no individual post-hoc comparison remained significant after Holm–Bonferroni correction. Thus, the statistical evidence supports an overall treatment-dependent spatial response, while substrate-specific differences in this response were not established (Table S1). These observations further emphasize that bulk rheology alone does not fully describe the mechanical environment near biological interfaces.

Taken together, our findings demonstrate that AC-ECM identity influences the spatial microrheological response of dSF, while StcE treatment produces a stronger normalized near-interface response across the investigated inter-faces. Although the spatial-viscosity increases more similarly following treatment, differences in the frequency-dependent spectra and relative elastic and viscous contributions persist. Given the established role of Lub’s mucin domain in cartilage lubrication and its susceptibility to StcE cleavage,[27] these findings are consistent with a potential contribution of lubricin to the spatial organization and mechanical response of SF.[1]

### 2.4 Limitations and Outlook

In this study, Col-II, Col-I, and FN were individually immobilized on glass as molecularly defined models of cartilage AC-ECM surfaces. This approach enables controlled comparisons of AC-ECM identity but does not reproduce the composition or structural organization of native articular cartilage, including the collagen-fibrillar network of its superficial zone.[42]

Interpretation using the generalized Stokes–Einstein framework also assumes that tracer motion samples a mechanically representative environment and that surface interactions are sufficiently controlled. Spatially heterogeneous glycoprotein-rich phases reported in HA–Lub model solutions reinforce the potential influence of the local microenvironment on tracer dynamics.[14, 43]

Although StcE has previously been shown to cleave Lub’s mucin domain in SF,[27] potential contributions from other StcE-sensitive glycoproteins cannot be excluded. While the frequency-dependent measurements revealed distinct elastic and viscous responses, including crossovers between *G*′ and *G*″ under some conditions, passive MPTM does not reproduce the coupled loading and sliding experienced during joint articulation. Given the dependence of cartilage mechanics on loading frequency and motion,[40, 41] the implications of these spatial and frequency-dependent responses for physiological lubrication remain to be established.

Future investigations could combine direct characterization of Lub cleavage with selective depletion and reconstitution experiments to establish its contribution to AC-ECM-dependent spatial microrheology. In particular, comparisons between intact and mucin-domain-modified recombinant Lub could help clarify how the mucinous domain of Lub influences the local mechanical response of SF. Given the growing interest in recombinant Lub as an intra-articular therapeutic lubricant,[44, 45] extending these measurements to native or engineered cartilage interfaces and correlating them with lubrication under mechanical loading could help establish their functional and therapeutic relevance. Previous surface force apparatus studies have shown that the contribution of HA-containing lubricant structures to friction and wear protection can depend on cartilage deformation and mechanical loading.[36]

We anticipate that our findings will assist mechanistic interpretation of direct tribological measurements of these biointerfaces in future work, and further motivate research on how molecularly distinct biological interfaces regulate the mechanical environment of adjacent complex fluids.

## 3 Conclusions

Depth-resolved MPTM revealed distinct spatial microrheological responses of SF near molecularly defined AC-ECM interfaces. Untreated dSF exhibited pronounced near-interface increases in apparent viscosity adjacent to Col-II and FN, whereas the response near Col-I was comparatively weak. StcE treatment substantially altered these responses, producing stronger and more similar normalized spatial-viscosity profiles across the three AC-ECM surface models. Statistical analysis further identified significant depth and fluid treatment × depth effects on the viscoelastic response. Comparison with QCM-D revealed that adsorption amount (at the nanoscale) did not directly predict the magnitude of spatial microrheological modulation for either of the SF-derived fluids. Together, these findings highlight the importance of AC-ECM molecular identity and mucin-domain integrity in shaping the tracer-scale mechanical response of SF near biological interfaces, consistent with a potential contribution from lubricin and other StcE-sensitive SF constituents.

## 4 Experimental Section

### 4.1 Materials

Phosphate-buffered saline (PBS; Gibco), absolute ethanol (200 proof, ≥99.5%; ACROS Organics), and ultrapure sodium dodecyl sulfate (SDS; ≥99%; MP Biomedicals) were used as received. (3-Aminopropyl)trimethoxysilane (APTMS; Thermo Scientific, USA, 97%) and glutaraldehyde solution (25% in water, Grade I; Sigma-Aldrich) were used for glass-surface functionalization. Cysteamine hydrochloride (Sigma-Aldrich, ≥98% titration) and glutaraldehyde were used for functionalization of gold-coated QCM-D crystals. Ultrapure water with a resistivity of 18.2 MΩ cm was obtained from a Thermo Fisher Scientific water-purification system. Bovine collagen type II (Col-II; SouthernBiotech), bovine collagen type I (Col-I; Gibco, Advanced Biomatrix), and fibronectin (FN; Invitrogen) were used as AC-ECM components. Bovine synovial fluid (SF; Lampire Biological Laboratories) was used as the lubricant preparation. Recombinant equine lubricin (rEqLub) was produced as described below. Carboxylated fluorescent tracer particles (Fluo-Spheres™, Invitrogen, Thermo Fisher Scientific, USA), with nominal diameters of 0.5 µm, red fluorescence, and excitation/emission wavelengths of 580/605 nm, were used for multiple-particle tracking microrheology (MPTM). Particle motion was recorded using an inverted epifluorescence microscope equipped with a TRITC fluorescence filter set.

All experiments involving biological materials were conducted in accordance with the procedures and guidelines of the Institutional Biosafety Committee at the University of California, Merced.

### 4.2 Synovial fluid preparation

Bovine SF was stored at −80 °C until use. On the day of each experiment, SF was thawed and centrifuged at 6000 rpm (~ 2000 *g*) for 5 min to remove cellular debris and large tissue aggregates. The centrifuged SF was then diluted to 25% (v/v) in PBS at pH 7.4. This preparation is referred to throughout the manuscript as diluted synovial fluid (dSF).

### 4.3 StcE treatment of synovial fluid

Centrifuged SF was treated with the mucin-selective met-alloprotease StcE-Δ35 to perturb densely O-glycosylated mucin-domain proteins. Recombinant StcE was expressed using a pET28b expression vector and purified by immobilized metal affinity chromatography.[46] SF was incubated with approximately 5.3 ng /mL StcE for 2 h at 37 °C under continuous stirring, following a previously reported protocol.[27] The treated SF was subsequently stored at −20 °C for at least 24 h. Before each experiment, samples were thawed and diluted to 25% (v/v) in PBS (pH 7.4). The resulting preparation is referred to as StcE-dSF.

### 4.4 Recombinant equine lubricin

Recombinant equine lubricin (rEqLub) was produced using a codon-scrambling strategy as previously described,[47] yielding a stock solution with a maximum concentration of 20 mg /mL. The stock solution was diluted to 5 mg/ mL in PBS and stored at −20 °C. For bulk rheology and MPTM measurements, aliquots were thawed and diluted to 250 µg/ mL in PBS on the day of each experiment.

### 4.5 Extracellular-matrix protein solutions

Col-II and Col-I were stored as received at 4 °C. Working solutions of each collagen were prepared at 50 µg /mL in PBS at pH 7.4 immediately before substrate functionalization. An FN solution (1 mg/mL in water) was stored at −20 °C, and working solutions of 50 µg /mL in PBS at pH 7.4 were prepared the day of use. For QCM-D measurements, this condition corresponds to the extended FN state characterized previously.[48]

### 4.6 Preparation of ECM-functionalized glass substrates

#### 4.6.1 APTMS/glutaraldehyde functionalization

Clean glass-bottom dishes (Cellvis, USA) with a dish diameter of 35 mm, glass-bottom well diameter of 10 mm and nomilar coverslip thickness of 0.17 mm, were functionalized with (3-aminopropyl)trimethoxysilane (APTMS) by applying APTMS to the glass surface using an APTMS-saturated cotton swab. The treated surfaces were allowed to react for 6 min at room temperature in a fume hood and were subsequently rinsed thoroughly with ultrapure water.

APTMS-functionalized surfaces were activated by depositing approximately 100 µL of 0.5% glutaraldehyde solution onto the treated region and incubating for 30 min at room temperature in the dark. Surfaces were subsequently rinsed thoroughly with ultrapure water before protein immobilization. Successful APTMS functionalization was independently confirmed by X-ray photoelectron spectroscopy (XPS), as described below.

#### 4.6.2 Immobilization of ECM proteins on glass

Following APTMS/glutaraldehyde functionalization, 100 µL of 50 µg /mL Col-II, Col-I, or FN in PBS was deposited onto the functionalized glass and incubated for 2 h at room temperature. Surfaces were subsequently rinsed with PBS to remove loosely bound molecules prior to use.

### 4.7 Depth-resolved multiple-particle tracking microrheology

#### 4.7.1 Sample preparation and image acquisition

Fluorescent carboxylated polystyrene tracer particles were added to dSF, StcE-treated dSF, or rEqLub at a 1:2000 (v/v) particle-suspension-to-sample ratio. The resulting suspensions were mixed and sonicated in a water bath for 30 s to disperse the tracer particles. Immediately following sonication, 100 µL of tracer-containing lubricant was deposited onto Col-II-, Col-I-, or FN-functionalized glass substrates and covered with a freshly cleaned coverslip. Samples were allowed to equilibrate for 45 min at room temperature in the dark before imaging.

Particle motion was imaged using an inverted epifluorescence microscope (Olympus IX 83 P2ZF) equipped with a 60× objective (Olympus LUCPlanFL N, numerical aperture 0.7). Videos were acquired at approximately 11 frames/ s with an image resolution of 2048 ×2048 pixels and a field of view of approximately 220 ×220 µm^2^, using a scientific video camera.

Measurements were acquired at defined distances from the ECM-fluid interface to quantify the spatial evolution of the local rheological response. Imaging planes were positioned at 0, 2, 4, 6, 8, 10, 12, 14, 16, 18, 20, 50, and 100 µm from the interface. The nominal *d* = 0 µm position was defined as the first imaging plane at which tracer particles were in focus; this position does not independently establish the particle-center-to-surface distance. For each condition, at least three independent samples were analyzed.

#### 4.7.2 Particle tracking and mean-squared displacement analysis

Particle positions and trajectories were tracked using Trackpy.[49] A static-localization-error baseline was determined using tracer particles immobilized within three independently prepared hydrogels with an elastic modulus of approximately 40 kPa, previously characterized by our group.[50] Five technical replicates were acquired for each hydrogel. The mean squared displacement (MSD) of each replicate was first averaged over the analyzed lag-time range, and the resulting five values were averaged to obtain a single baseline estimate for each hydrogel. The mean of the three hydrogel-level estimates was then subtracted from each experimental MSD as a lag-time-independent offset. The resulting static-localization-error baseline was 0.0022± 0.0002 µm^2^ (mean ±SEM; *n* = 3 independently prepared hydrogels). This correction had a negligible effect on the calculated apparent viscosities, changing the reported values only at the third decimal place.

For each imaging depth, the two-dimensional mean-squared displacement (MSD) was calculated as a function of lag time, τ, according to

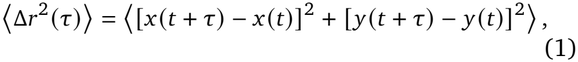

where *x* and *y* denote the in-plane particle coordinates. The time-averaged MSD was first calculated for each particle trajectory using all available time origins, followed by ensemble averaging across particles.

For normal Brownian motion in a homogeneous Newtonian fluid, the two-dimensional MSD increases linearly with lag time, ⟨Δ*r*^2^ (τ)⟩ = 4*D*τ, where *D* is the translational diffusion coefficient. Deviations from this linear scaling are characteristic of anomalous diffusion, which can be described by a power-law dependence on lag time. Following established particle-tracking analyses of complex biological fluids,[31, 51] each experimental MSD was fitted over a selected lag-time interval (τ≈ 0.09– 4.5 s) using

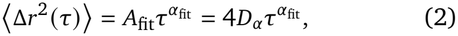

where *D*_α_ is the generalized diffusion coefficient and α_fit_ is the anomalous diffusion exponent. Normal Brownian diffusion corresponds to α_fit_ = 1, whereas α_fit_ < 1 indicates subdiffusive motion over the fitted interval. The resulting parameters characterize the overall scaling of the ensemble-averaged MSD. A representative workflow of the particle-tracking analysis performed using Trackpy is provided in SI (Section S7).

#### 4.7.3 Calculation of apparent viscosities

For anomalous diffusion, the generalized diffusion coefficient *D*_α_ has dimensions of length squared per time raised to α_fit_, and therefore differs from the conventional diffusion coefficient of Brownian motion. To compare tracer mobility across conditions with different anomalous diffusion exponents, an effective diffusivity was defined at a fixed reference lag time, τ_0_, as

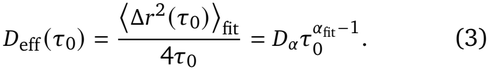

A reference lag time of τ_0_ = 1 s was selected for all conditions, consistent with previous particle-tracking studies employing fixed-timescale effective diffusivities to characterize particle mobility in complex biological fluids.[51, 52] The corresponding apparent viscosity was estimated using the Stokes–Einstein relation,[53]

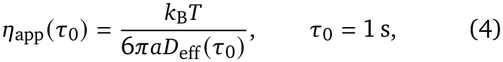

where *k*_B_ is the Boltzmann constant, *T* = 298.15 K is the absolute temperature, and *a* is the tracer-particle radius. For subdiffusive motion, *D*_eff_ (τ_0_) represents a timescale-dependent measure of particle mobility rather than a constant Fickian diffusion coefficient. Accordingly, η_app_ (τ_0_) is interpreted as a tracer-mobility-based apparent viscosity at the selected reference timescale, rather than an intrinsic steady-shear viscosity. Hydrodynamic interactions near solid boundaries may additionally influence tracer mobility.[35]

The global power-law fit was used to characterize the overall MSD scaling and calculate this scalar apparent-viscosity estimate. Frequency-dependent viscoelastic properties were obtained separately from the MSD using the generalized Stokes–Einstein framework and local logarithmic scaling exponents, as described below.[28, 31]

To compare depth-dependent variations across extra-cellular matrix (ECM) surfaces and fluid conditions, the apparent viscosity was normalized using a condition-specific far-field apparent viscosity reference. For each experimental condition, the apparent viscosities obtained from independent measurements were first averaged separately at *d* = 50 µm and *d* = 100 µm. The reference was defined as the mean of these two depth-specific averages,

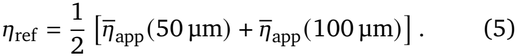

The same reference was applied to the individual independent measurements of that condition to obtain

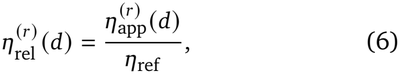

where *r* identifies an independent measurement. The normalized values were subsequently averaged at each depth and reported as mean ±standard error of the mean (SEM) across independent measurements. This procedure places the far-field reference region near unity without requiring identical apparent viscosities at 50 µm and 100 µm individually.

#### 4.7.4 Conversion of MSDs to linear viscoelastic properties

For the frequency-dependent microrheological analysis, the MSD was related to the shear creep compliance, *J* (τ), using the generalized Stokes–Einstein framework.[28] For the two-dimensional MSD measured here,

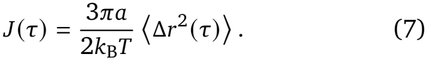

The creep compliance was converted to the complex modulus and complex viscosity according to established procedures.[28, 43] Whereas the global power-law fit provides a single scaling exponent over the selected lag-time interval, the frequency-dependent analysis uses the local logarithmic slope of the MSD,

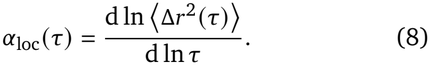

α_loc_ (τ) was estimated from local power-law fits using a centered, seven-point moving window, with up to three neighboring lag-time points on either side.[43] Unlike the global exponent α_fit_, the local exponent can vary with lag time, allowing changes in the MSD scaling behavior to be represented across the observation window.

Following the local power-law approximation, angular frequency was associated with lag time according to

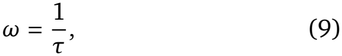

and the frequency-dependent local exponent was defined as

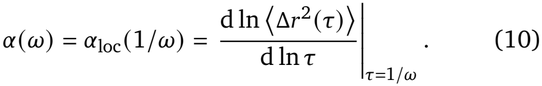

Thus, α ω represents the same local MSD scaling exponent evaluated at the lag time corresponding to angular frequency ω. The frequency-dependent complex shear modulus, *G*\* (ω) = *G*′ (ω)+ *iG*″ (ω), was estimated using the local power-law approximation,[43]

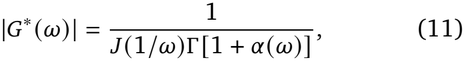

where Γ is the gamma function. The storage and loss moduli were then obtained from

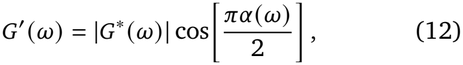

and

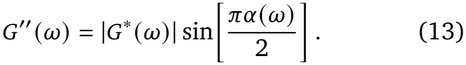

The magnitude of the complex viscosity was calculated as

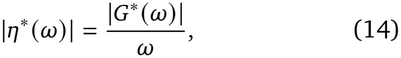

The apparent viscosity η_app_ and the complex-viscosity magnitude |η* (ω)| represent distinct quantities and were evaluated separately, although for many fluids their magnitudes are approximately equal.[54]

#### 4.7.5 Constitutive modeling of linear viscoelastic properties

The frequency-dependent moduli were fitted with six candidate linear viscoelastic constitutive models based on combinations of classical and fractional rheological elements:[55, 56] (i) a Newtonian fluid, (ii) a springpot, (iii) a fractional Kelvin–Voigt liquid consisting of a springpot in parallel with a Newtonian viscous contribution,[57] (iv) a classical Jeffreys model consisting of a Maxwell element in parallel with a Newtonian viscous contribution, (v) a fractional Maxwell liquid, and (vi) a fractional Jeffreys-type model consisting of a fractional Maxwell liquid branch in parallel with a Newtonian viscous contribution. In the following expressions, *s* = *i*ω.

The six models are

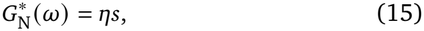

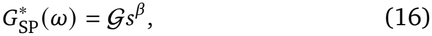

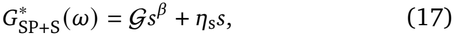

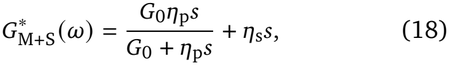

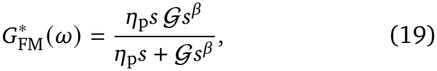

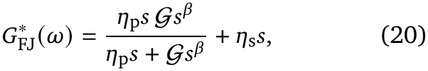

respectively. η is the viscosity of a Newtonian fluid; η_p_ is the dashpot viscosity within a series branch; η_s_ is the effective viscosity of a parallel Newtonian contribution; *G*_0_ is the elastic modulus of the classical Maxwell spring; and *G* is the springpot coefficient, with units of Pa s^β^. The fractional exponent β is dimensionless, with 0 < β < 1, and controls the intermediate power-law response between ideal elastic and viscous behavior. The term η_s_ is an effective constitutive parameter and was not independently assumed to equal the viscosity of the aqueous solvent.

The most general two-element models are the fractional Maxwell and the fractional Kelvin-Voigt, each containing two spring-pots, either in series or in parallel. For the fractional Kelvin–Voigt liquid, fractional Maxwell liquid, and fractional Jeffreys-type models, one fractional exponent was fixed to unity, reducing the corresponding element to a Newtonian dashpot while retaining a single fractional exponent, β. This restriction has been employed in fractional constitutive descriptions of mucin-rich systems and viscoelastic polymer solutions.[57, 58] The fractional Jeffreys-type model employed here extends the fractional Maxwell liquid by adding a parallel Newtonian viscous contribution, following the general fractional Jeffreys architecture that has been employed for complex biopolymer fluids.[59] A closely related fractional Cross representation has also been applied to HA– rhPRG4 mixtures.[14] The parameter values are used here descriptively and are not interpreted as uniquely determined molecular relaxation mechanisms.

For each measurement condition and depth, experimental *G*′ and *G*″ spectra from independent measurements were aligned by angular frequency and averaged. Model parameters were obtained by simultaneously fitting the two mean moduli over 0.2 rad/s≤ ω ≤12 rad/s using bounded nonlinear least squares (`scipy.optimize.least_squares`).[60] For each candidate model, parameter estimation minimized the combined residual objective, 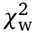, computed by

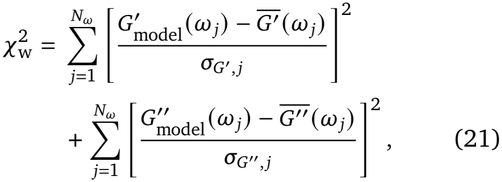

where 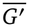 and 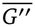 are the replicate-averaged moduli, *N*_ω_ is the number of analyzed frequencies, and 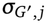 and 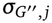 are the corresponding SEMs. To avoid excessive weighting of data points with very small SEMs, a minimum SEM was set at 10% of the median nonzero SEM for each modulus across the fitted frequency range. Positive parameters were fitted on logarithmic parameter scales, whereas β was bounded to 0.01 ≤β≤ 0.99. The modeled complex viscosity was calculated directly from the fitted complex modulus, 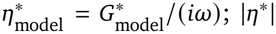 was not treated as an additional independent fitting target.

Candidate models were ranked by an approximate small-sample-corrected Akaike information criterion (AICc), computed consistently from the weighted residual objective as

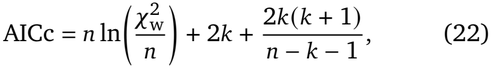

where *n* = 2*N*_ω_ is the number of fitted modulus values and *k* is the number of fitted constitutive parameters. The candidate with the lowest AICc was identified as the preferred descriptive model, provided its fit was numerically stable and residuals did not exhibit clear systematic structure. Because the frequency-domain moduli are both derived from the same particle trajectories, residuals may be correlated across frequencies and between moduli; moreover, the SEMs were estimated from a limited number of independent measurements. Accordingly, these AICc values were used as approximate comparative scores, not as formal probabilities that a particular constitutive model is physically correct. From 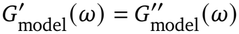, crossovers were identified within the fitted frequency interval.

The depth-dependent normalized apparent viscosity was additionally analyzed using single- and double-Boltzmann sigmoidal decay functions. These empirical functions describe the transition between increased near-interface apparent viscosity and a far-field reference value, without imposing a microscopic model for the origin of the spatial gradient. The single-Boltzmann model was

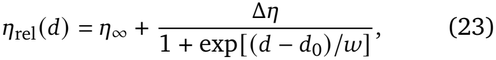

where η_∞_ is the far-field relative-viscosity asymptote, Δη > 0 is the fitted increased apparent viscosity, *d*_0_ is the midpoint of the sigmoidal transition, and *w* > 0 is its characteristic width. A second model allowed two transitions with potentially distinct midpoints and widths,

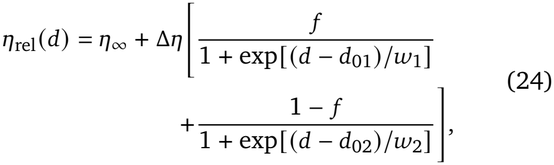

where 0 < *f* < 1 partitions the total fitted increase between two decaying contributions, *d*_01_ < *d*_02_ are their respective transition midpoints, and *w*_1_, *w*_2_ > 0 are their widths. The parameters Δη and *f* do not directly quantify separate molecular populations or film masses. In the implemented comparison, η_∞_ was fixed to unity, consistent with the far-field normalization, leaving three adjustable parameters for the single-Boltzmann model and six for the double-Boltzmann model.

Fits were performed on the replicate-averaged η_rel_ (*d*) profiles over 0 µm≤ *d* ≤20 µm, while the measurements at 50 µm and 100 µm defined the reference used for normalization. Both decay functions were optimized by bounded nonlinear least squares (`scipy.optimize.curve_fit`),[60] using the SEM of the normalized viscosity as the weighting uncertainty. A small uncertainty floor, defined as the larger of 0.025 relative-viscosity units or 10% of the median positive SEM, was applied during fitting to prevent near-zero SEMs from dominating the optimization. Decay amplitudes and widths were constrained to positive values. The widths were restricted to at least 0.5 µm to discourage unresolved step-like transitions, and the two midpoints in the double-Boltzmann model were ordered to prevent component interchange. The two spatial-decay models were compared using Eq. (22), with *n* replaced by the number of fitted depth-averaged values, *N*_*d*_, *k* = 3 or 6, respectively, and 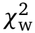 replaced by the SEM-weighted spatial residual sum of squares. Model selection was considered only when the number of fitted depths was sufficient for the AICc correction to be defined. The change in AICc, the fitted residual patterns, convergence status, and parameter identifiability were assessed together when evaluating whether the additional flexibility of the double-Boltzmann function was justified. To quantify the spatial extent of the apparent-viscosity enhancement, a characteristic distance *d*_95_ was defined as the distance at which the fitted excess apparent viscosity decreased to 5% of the total increased amplitude relative to 1. For single-sigmoid fits, this amplitude corresponds to (*A*), whereas for double-sigmoid fits, it corresponds to (*A*_1_+ *A*_2_).

Data processing, rheological calculations, constitutive modeling, and spatial-decay analysis were performed using custom Python scripts with NumPy, SciPy, and pandas.[61]

### 4.8 MPTM statistical analysis

Statistical analyses were performed at the level of independent experimental preparations. For MPTM, *G*′, *G*″, and |η*| were evaluated directly at ω = 1 rad/s for each independent measurement, without constitutive-model fitting. The AC-ECM analysis included dSF and StcE-dSF measured at 2 and 100 µm from Col-II, Col-I, and FN surfaces. A three-factor split-plot ANOVA was used to evaluate fluid treatment, ECM identity, depth, and their interactions, accounting for repeated depth measurements within the same preparation. For the two measured depths, the repeated-measures design was implemented using separate Type-III, sum-coded factorial models of each preparation’s depth-averaged response and paired depth difference.

When a relevant ANOVA interaction was significant, post-hoc comparisons were performed using paired *t*-tests for within-experiment depth differences and Welch’s *t*-tests for comparisons between independent preparations. Resulting *p*-values were adjusted with Holm-Bonferroni correction. Far-field MPTM measurements on glass at 100 µm were analyzed separately using one-way ANOVA to compare dSF, StcE-dSF, and recombinant equine lubricin (rEqLub). When the ANOVA test was significant, pairwise Welch’s *t*-tests with Holm–Bonferroni correction were performed.

For factorial ANOVAs, heteroscedasticity-robust HC3 *p*-values were additionally calculated as sensitivity analyses. Statistical significance was defined as *p* < 0.05 for ANOVA tests and Holm-adjusted *p* < 0.05 for post-hoc comparisons. Multiplicity corrections were applied within the specified comparison families rather than across all ANOVA effects or rheological quantities. Reported confidence intervals for individual contrasts were unadjusted 95% intervals. Analyses were performed using custom Python scripts with NumPy, SciPy, and statsmodels.[60, 62, 63]

### 4.9 Complementary characterization techniques

Quartz crystal microbalance with dissipation (QCM-D), atomic force microscopy (AFM), and X-ray photoelectron spectroscopy (XPS) were used as complementary techniques to characterize StcE-dSF adsorption on AC-ECM models, FN morphology and nanomechanics, and glass functionalization, respectively. Methods details along with relevant methodological references for QCM-D,[21, 22, 64, 65] AFM,[21, 22, 66–70] and XPS[71] are found in Supplementary Information, Sections S1, S2, and S6.

## Author Contributions

**D.R.J.P**.: Conceptualization, Methodology, Investigation, Formal analysis, Visualization, Data curation, Software, Validation, Supervision, Writing – original draft, Writing – review & editing.

**N.C.R**.: Investigation, Formal analysis, Visualization, Data curation, Software, Writing – original draft, Writing – review & editing.

**N.L.M**.: Investigation, Writing – review & editing. **M.J.P**.: Resources, Investigation, Funding acquisition, Writing – review & editing.

**H.L.R**.: Resources, Investigation, Funding acquisition, Writing – review & editing.

**L.J.B**.: Resources, Funding acquisition, Methodology, Writing – review & editing.

**G.D.D**.: Methodology, Validation, Supervision, Writing – review & editing.

**R.C.A.E**.: Conceptualization, Methodology, Visualization, Validation, Supervision, Funding acquisition, Project administration, Resources, Writing – review & editing.

## Data Availability Statement

The data that support the findings of this study are available from the corresponding author upon reasonable request.

## Acknowledgments

D.R.J.P., N.C.R., and R.C.A.E. acknowledge funding from the National Science Foundation (NSF) CAREER Award through NSF-CMMI-2239665 awarded to R.C.A.E, and UC Merced Startup funds awarded to R.C.A.E. R.C.A.E., D.R.J.P., N.L.M., M.J.P, H.L.R. and L.J.B. acknowledge NSF LEAP HI Award funding through NSF-CMMI-2245367 awarded to M.J.P., H.L.R., L.J.B. and R.C.A.E.

We acknowledge funding from the NSF-CREST Center for Cellular and Biomolecular Machines (CCBM) at UC Merced (NSF-HRD-1547848). N.C.R. acknowledges the funding from CCBM fellowship through NSF-HRD-1547848. D.R.J.P. and R.C.A.E. thank Dr. Mourad Sadqi for his continued technical support. D.R.J.P. and N.C.R. thank Surbhi Salgarkar for help with hydrogel preparation for static video error calibration. D.R.J.P. thanks Sixin Zhai for her support and helpful feedback throughout this project, as well as Professors Christopher Viney and Linda Hirst for their valuable input as members of his doctoral committee.

