## Supplemental Materials for "Synovial Fluid Spatial Microrheology is Shaped by Extracellular Matrix Identity and Mucin Integrity"

### List of Supplementary Figures and Tables

| | | |
|---|---|---|
| Figure S1 | High-resolution N 1s XPS spectra of APTMS-functionalized and untreated glass. | 2 |
| Figure S2 | AFM characterization of the morphology and apparent nanomechanical properties of immobilized FN. | 3 |
| Figure S3 | Depth-dependent viscosity profiles for dSF on surfaces and Boltzmann fitting diagnostics. | 4 |
| Figure S4 | Depth-dependent viscosity profiles for StcE-dSF on surfaces and Boltzmann fitting diagnostics. | 5 |
| Figure S5 | Depth-dependent viscosity profiles for rEqLub on surfaces and Boltzmann fitting diagnostics. | 6 |
| Figure S6 | Depth-dependent viscosity profiles for rEqLub on surfaces and Boltzmann fitting diagnostics. | 7 |
| Table S1 | Effects of fluid treatment and measurement depth on synovial fluid microrheology. | 8 |
| Figure S7 | QCM-D raw traces of StcE-dSF assembly on Col-II, Col-I, and FN surfaces. | 11 |
| Figure S8 | Statistical analysis of synovial fluid adsorption and retention onto ECM substrates. | 12 |
| Figure S9 | Analysis workflow of tracer-particles using Trackpy. | 13 |
| Figure S10 | Representative drift-corrected tracer-particle trajectories in dSF and PBS. | 13 |

### S1 Glass functionalization with APTMS

A Thermo Scientific Nexsa G2 XPS system was used to confirm functionalization of glass substrates with APTMS. Glass substrates were prepared according to the procedure described above up to, but excluding, the glutaraldehyde coupling step. XPS data were analyzed using KherveFitting version 1.81.[1] Binding energies were charge-corrected by referencing the C-C component of the C 1s spectrum to 284.8 eV. For each sample, 25 scans were acquired at a single location near the center of the substrate using an X-ray spot size of 50 µm. Survey spectra were acquired using a 1 eV step size, a 10 ms dwell time, and a 200 eV pass energy. High-resolution spectra of the C 1s, N 1s, O 1s, and Si 2p regions were acquired using a 0.1 eV step size, a 50 ms dwell time, and a 50 eV pass energy. N 1s spectra of APTMS-functionalized substrates were compared with those of untreated glass to assess surface functionalization. High-resolution spectra were fitted using Gaussian-Lorentzian line shapes.

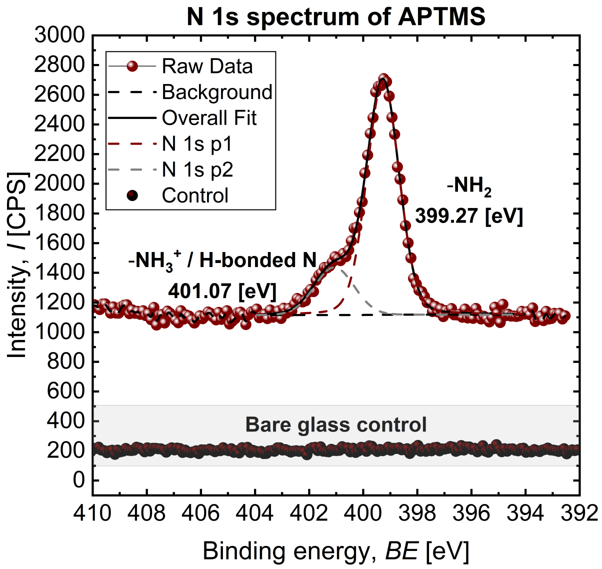


**Figure S1.** High-resolution N 1s XPS spectra of APTMS-functionalized glass and untreated glass. The APTMS-functionalized surface exhibits a clear N 1s signal that was fitted with two components centered at 399.27 and 401.07 eV, assigned to neutral amine (–$NH_2$) and protonated and/or hydrogen-bonded nitrogen (–$NH_3^+$/H-bonded N), respectively. No detectable N 1s signal was observed for untreated glass.

### S2 AFM characterization of FN

Atomic force microscopy (AFM) was used to characterize the surface morphology and apparent normal nanomechanics of immobilized fibronectin (FN) films. Gold-coated QCM-D crystals were functionalized with cysteamine and glutaraldehyde following the procedure described for QCM-D measurements. Following glutaraldehyde activation, the surfaces were incubated with 50 µg mL$^{-1}$ FN using the previously reported immobilization procedure.[2] Measurements were performed in PBS using a NanoWizard Pure AFM system (Bruker, USA) equipped with V-shaped ScanAsyst-Fluid probes (Bruker, USA) with gold-coated backsides and a nominal normal spring constant of 0.7 N m$^{-1}$. Surface topography was acquired in Quantitative Imaging (QI) mode using a controlled tip–sample force setpoint to minimize deformation of the hydrated protein films. Height images were processed using Gwyddion version 2.7.1.[3] Second-order polynomial row alignment and horizontal scar correction were applied before calculating the root-mean-square roughness, $S_q$, from the AFM height map. The representative FN-coated region shown in Supporting Information (Figure S3) exhibited an $S_q$ of 2.2 nm. Spatially resolved nanomechanical measurements were performed in three distinct regions of the FN-coated surface, each covering an area of $2 \times 2\ \mu m^2$. Each force-volume map consisted of 16,384 force–distance curves. The approach segments were fitted using the Hertz contact model in JPKSPM Data Processing software (version 8.1.62) to estimate the apparent Young's modulus, $E$.[4, 5] For each curve, the fit was restricted to 4–20% of the approach segment, with 0% corresponding to its final point and 100% to its initial point. This fitting interval was selected to limit potential contributions from the underlying substrate. A Poisson's ratio of $\nu = 0.5$ was assumed as a first-order approximation for the hydrated protein films. The extracted moduli were interpreted as apparent values because Hertz-model estimates are subject to uncertainties in the effective tip–sample contact geometry, the assumed Poisson's ratio, and potential contributions from the underlying substrate.[6, 7] Modulus distributions were represented as probability-density histograms using a bin width of 0.33 MPa. Measurements from the three regions were presented individually and as a pooled distribution.

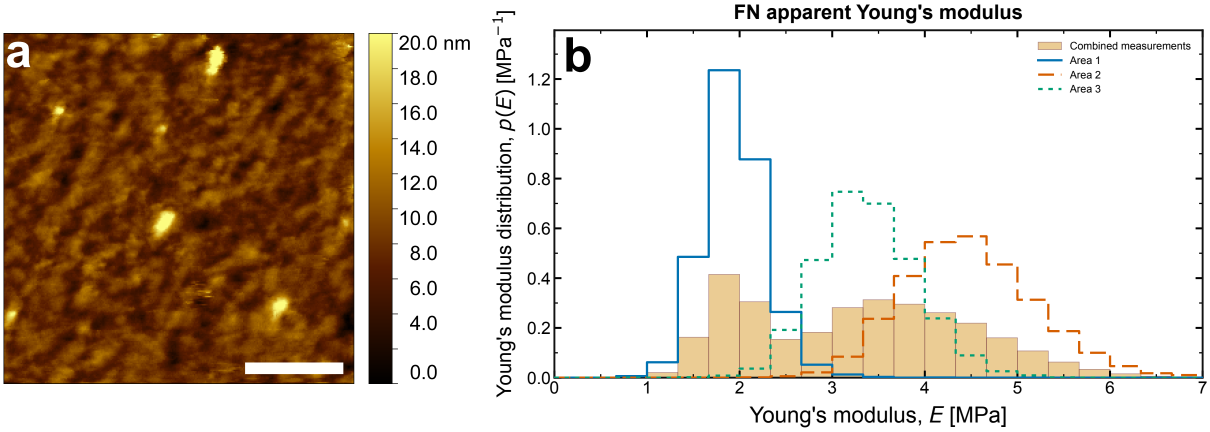


**Figure S2.** AFM characterization of immobilized fibronectin (FN) on gold. **a,** Representative AFM topography image of the FN-functionalized surface, exhibiting an areal root-mean-square roughness of $S_q = 2.2$ nm. Scale bar is 500 nm. **b,** Probability density distributions of apparent Young's modulus, $E$, obtained by AFM nanoindentation across three spatially distinct regions. The filled histogram represents the pooled measurements, while outlined histograms show the individual regions. All distributions are normalized to probability density.

### S3 Apparent viscosity controls and fit information

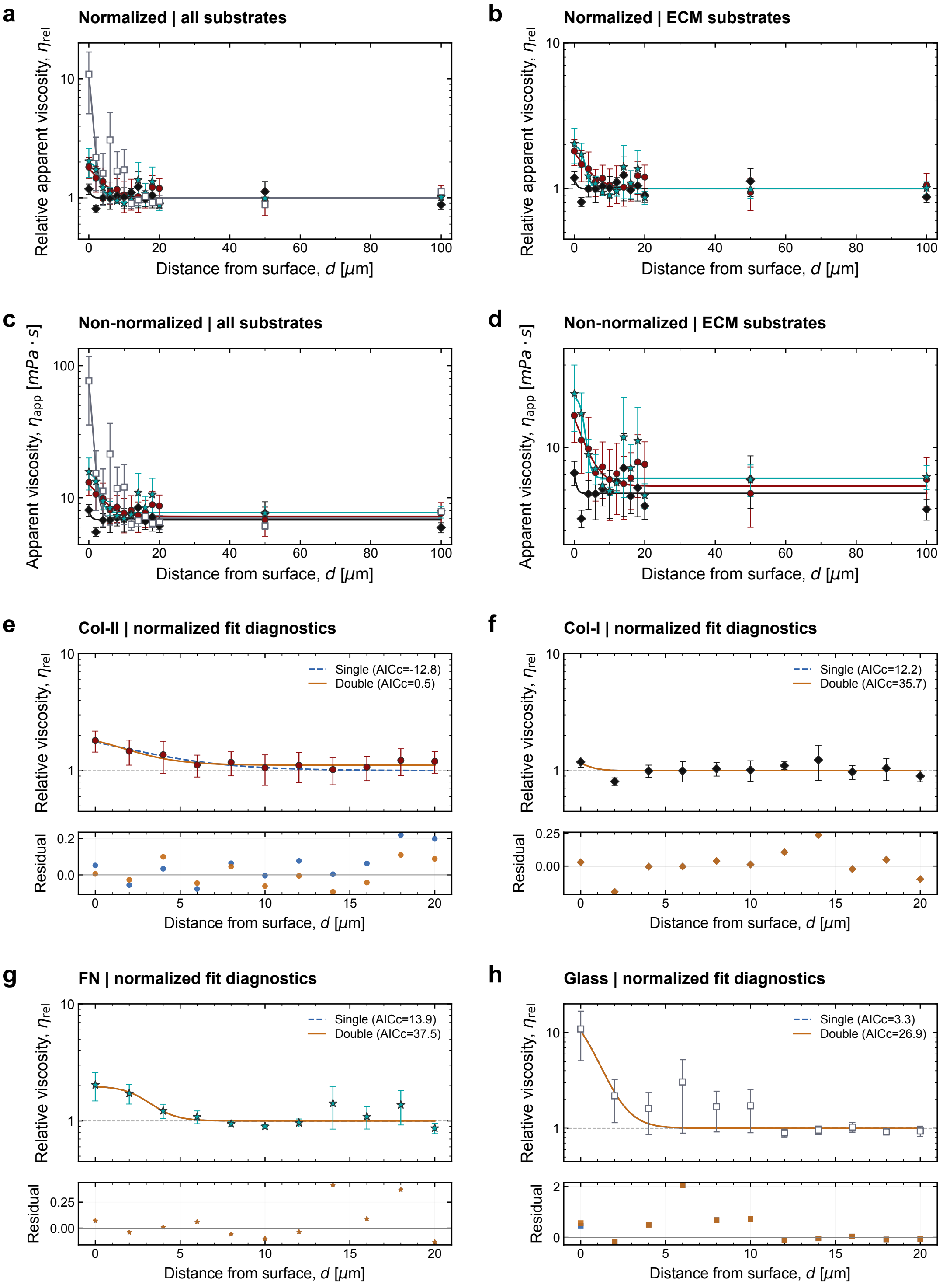


**Figure S3. a,b,** Relative apparent viscosity of dSF, $\eta_{\mathrm{rel}}$, as a function of distance from immobilized Col-II, Col-I, FN, and glass surfaces, shown for all substrates (a) and ECM substrates only (b). **c,d,** Corresponding non-normalized apparent viscosity profiles (mPa·s) for all substrates (c) and ECM substrates only (d). **e–h,** Comparison of single- and double-Boltzmann models fitted to normalized viscosity profiles within 0–20 $\mu$m of Col-II (e), Col-I (f), FN (g), and glass (h) surfaces. Residuals, calculated as the difference between experimental mean values and model predictions, are shown below each fit. Symbols represent mean values, and error bars indicate the standard error of the mean (SEM) across independent runs.

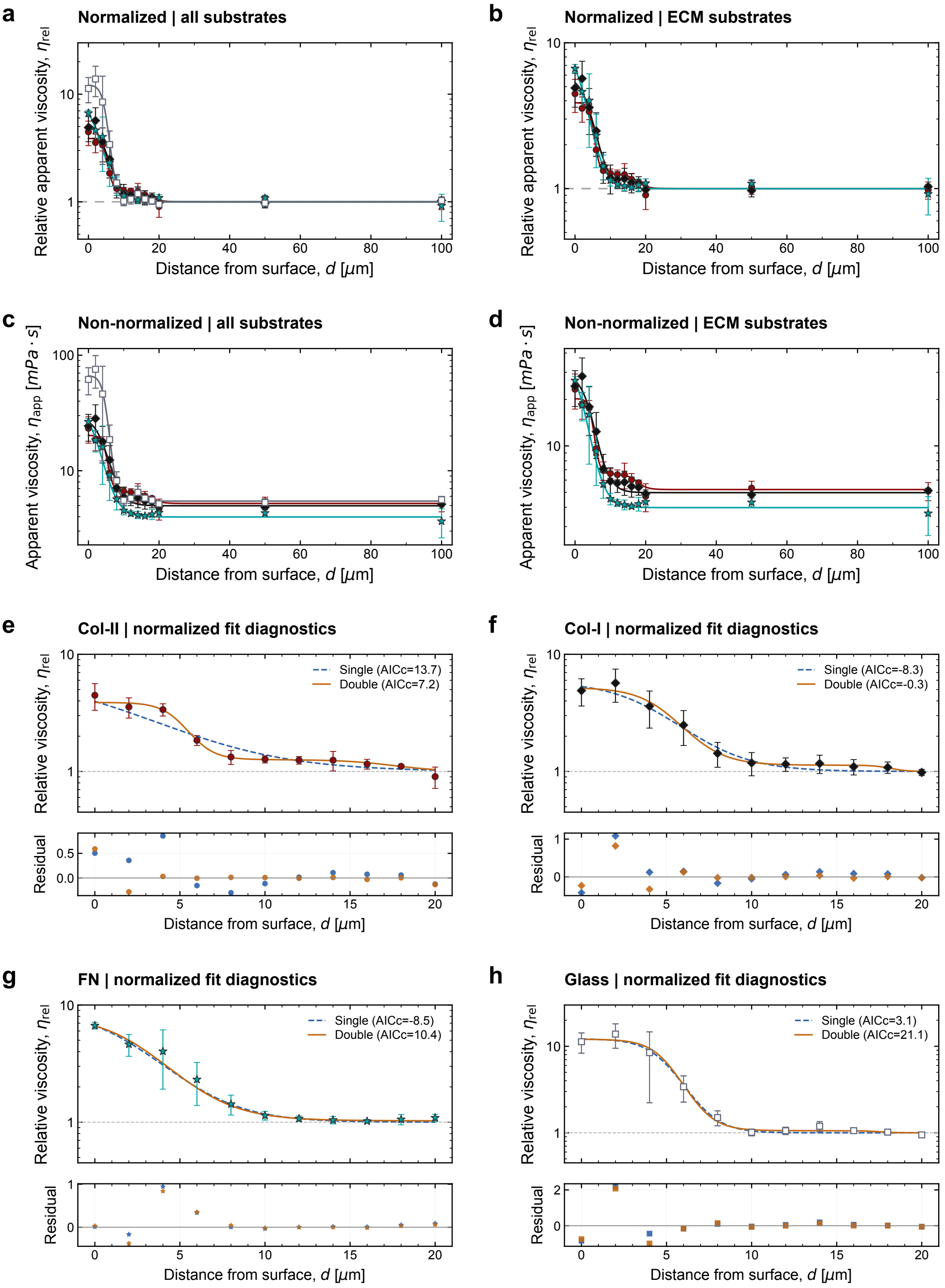


**Figure S4.** **a,b,** Relative apparent viscosity of Stce-dSF, $\eta_{\mathrm{rel}}$, as a function of distance from immobilized Col-II, Col-I, FN, and glass surfaces, shown for all substrates (a) and ECM substrates only (b). **c,d,** Corresponding non-normalized apparent viscosity profiles (mPa·s) for all substrates (c) and ECM substrates only (d). **e–h,** Comparison of single- and double-Boltzmann models fitted to normalized viscosity profiles within 0–20 $\mu$m of Col-II (e), Col-I (f), FN (g), and glass (h) surfaces. Residuals, calculated as the difference between experimental mean values and model predictions, are shown below each fit. Symbols represent mean values, and error bars indicate the standard error of the mean (SEM) across independent runs.

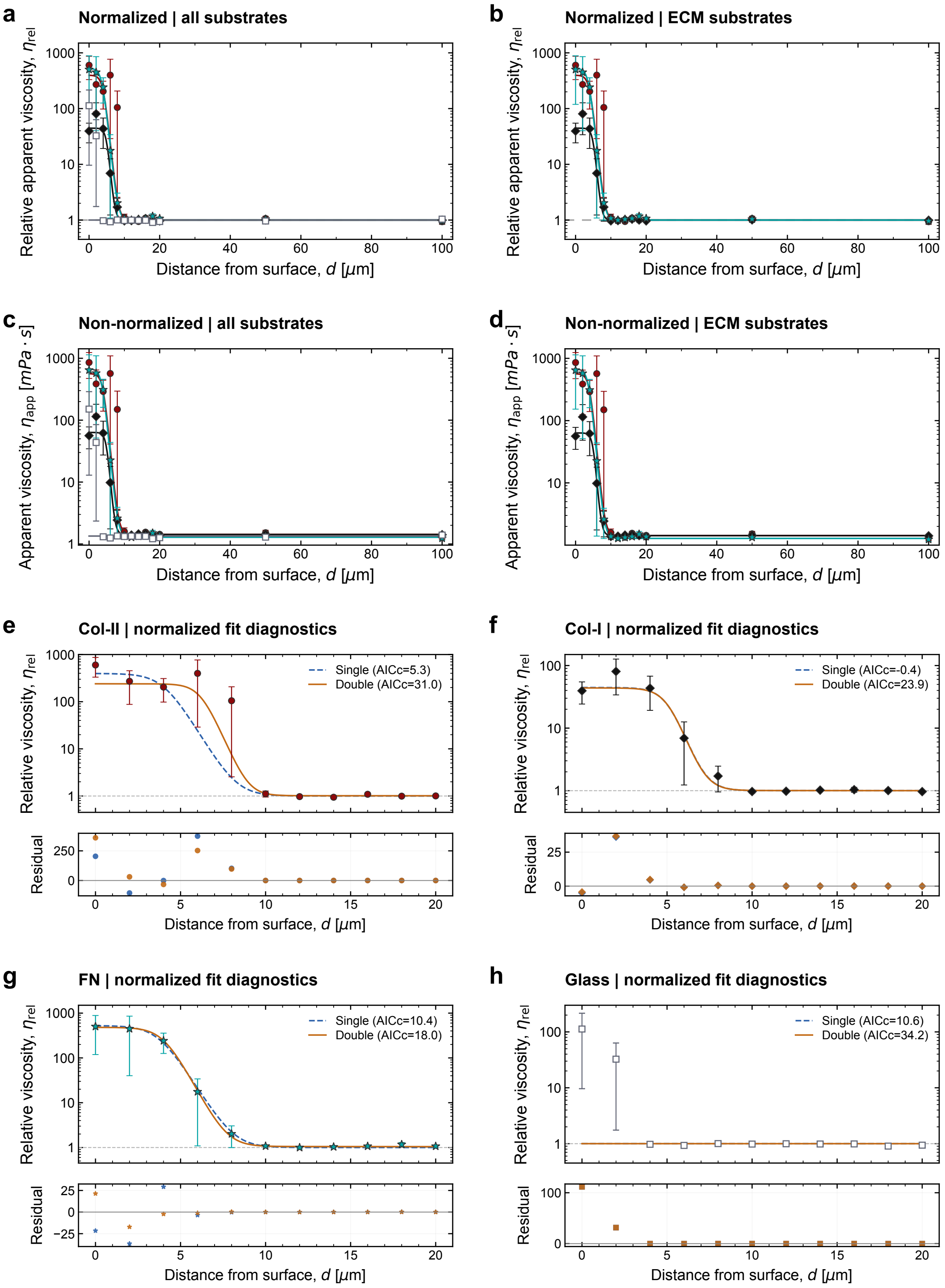


**Figure S5.** **a,b,** Relative apparent viscosity, $\eta_{\mathrm{rel}}$, of $250\,\mu\mathrm{g\,mL^{-1}}$ rEqLub in PBS as a function of distance from immobilized Col-II, Col-I, FN, and glass surfaces, shown for all substrates (a) and ECM substrates only (b). **c,d,** Corresponding non-normalized apparent viscosity profiles (mPa·s) for all substrates (c) and ECM substrates only (d). **e–h,** Comparison of single- and double-Boltzmann models fitted to normalized viscosity profiles within 0–20 $\mu$m of Col-II (e), Col-I (f), FN (g), and glass (h) surfaces. Residuals, calculated as the difference between experimental mean values and model predictions, are shown below each fit. Symbols represent mean values, and error bars indicate the standard error of the mean (SEM) across independent runs.

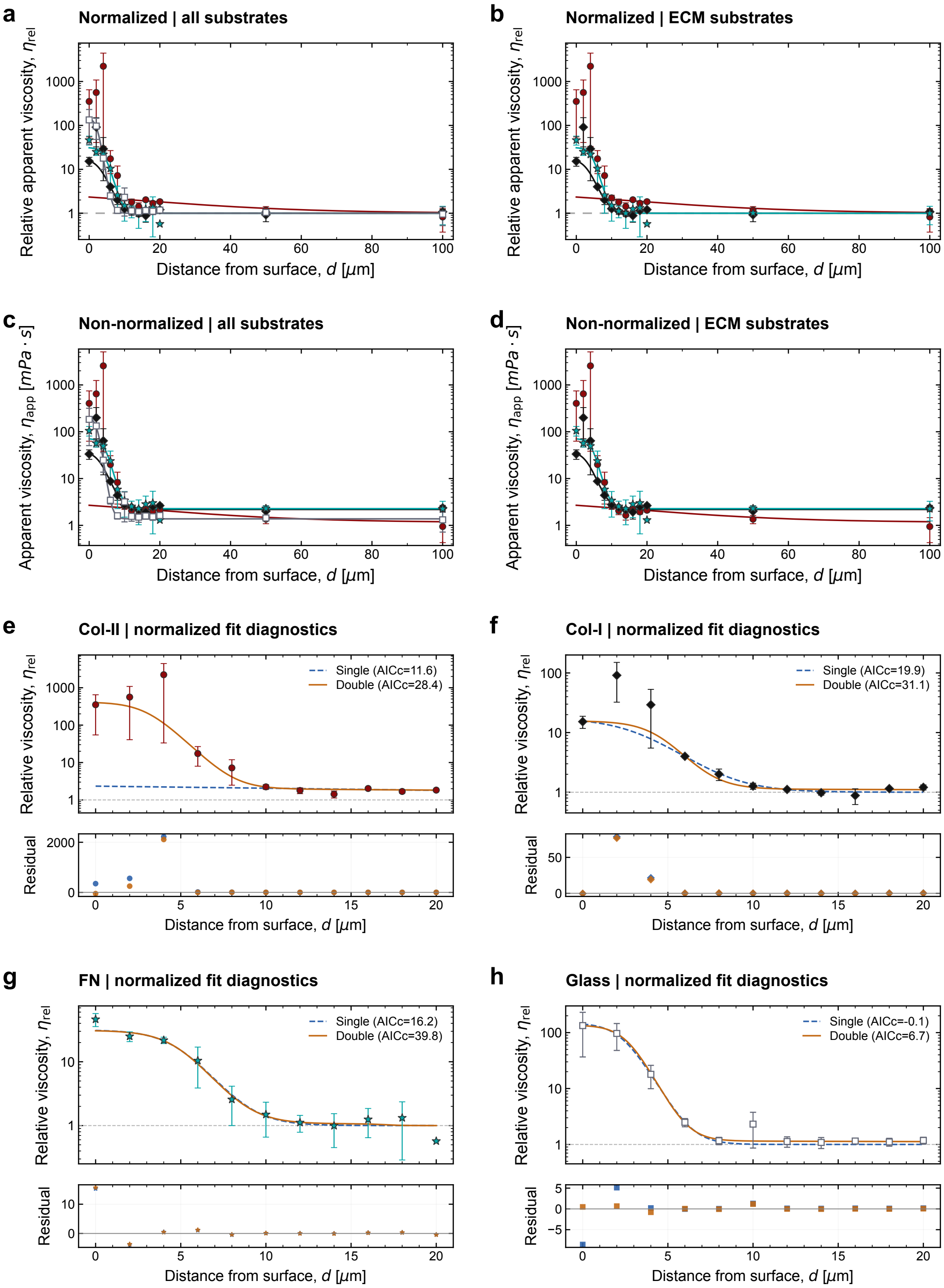


**Figure S6. a,b,** Relative apparent viscosity, $\eta_{rel}$, of PBS as a function of distance from immobilized Col-II, Col-I, FN, and glass surfaces, shown for all substrates (a) and ECM substrates only (b). **c,d,** Corresponding non-normalized apparent viscosity profiles (mPa·s) for all substrates (c) and ECM substrates only (d). **e–h,** Comparison of single- and double-Boltzmann models fitted to normalized viscosity profiles within 0–20 $\mu$m of Col-II (e), Col-I (f), FN (g), and glass (h) surfaces. Residuals, calculated as the difference between experimental mean values and model predictions, are shown below each fit. Symbols represent mean values, and error bars indicate the standard error of the mean (SEM) across independent runs.

### S4 Estimation of hydrodynamic wall effects on tracer particles

Following Faxén's approximation for the hydrodynamic drag of a spherical tracer translating parallel to a planar, no-slip boundary,

$$\frac{\gamma_{\parallel}}{\gamma_0} = \left[1 - \frac{9R}{16h} + \frac{R^3}{8h^3} - \frac{45R^4}{256h^4} - \frac{R^5}{16h^5}\right]^{-1} \approx \frac{1}{0.93} \approx 1.075, \tag{S1}$$

considering $R = 0.25\,\mu\text{m}$ and $h = 2\,\mu\text{m}$. This means that at 2 µm from the interface, the wall effect increases the parallel drag by approximately 7.5 %, which might lead to an error of approximately 7.5 % in the measured rheological properties. At $h = 100\,\mu\text{m}$, this effect is negligible (approximately 0.1 %).

### S5 ANOVA comparisons for viscoelastic properties of dSFs at 1 rad/s

**Table S1. Effects of fluid treatment and measurement depth on synovial fluid microrheology.** Selected results from a three-factor split-plot ANOVA of storage modulus ($G'$), loss modulus ($G''$), and complex-viscosity magnitude ($|\eta^*|$) at $\omega = 1\ \text{rad s}^{-1}$. Fluid treatment (dSF or StcE-dSF) and AC-ECM identity (Col-II, Col-I, or FN) were between-preparation factors, whereas measurement depth (2 or 100 µm) was a within-preparation factor. The $F$ statistic reports the conventional ANOVA test, with numerator and denominator degrees of freedom in parentheses. $p$ (OLS) denotes the conventional ANOVA $p$-value, and $p$ (HC3) denotes the heteroscedasticity-robust $p$-value calculated using the HC3 covariance estimator. Partial $\eta^2$ represents the proportion of effect-plus-error variance attributable to each effect, and $N$ is the number of independent experimental preparations. Only the depth and fluid treatment × depth effects are displayed.

| **Response** | **Effect** | ***F* statistic** | ***p* (OLS)** | ***p* (HC3)** | **Partial $\eta^2$** | $N$ |
|---|---|---|---|---|---|---|
| $G'$ | Depth | $F(1,13) = 22.76$ | $3.65 \times 10^{-4}$ | 0.00282 | 0.636 | 19 |
| $G'$ | Fluid × Depth | $F(1,13) = 11.70$ | 0.00455 | 0.02069 | 0.474 | 19 |
| $G''$ | Depth | $F(1,13) = 49.17$ | $9.17 \times 10^{-6}$ | $1.13 \times 10^{-4}$ | 0.791 | 19 |
| $G''$ | Fluid × Depth | $F(1,13) = 8.18$ | 0.01338 | 0.04475 | 0.386 | 19 |
| $\lvert\eta^*\rvert$ | Depth | $F(1,13) = 39.20$ | $2.92 \times 10^{-5}$ | $3.28 \times 10^{-4}$ | 0.751 | 19 |
| $\lvert\eta^*\rvert$ | Fluid × Depth | $F(1,13) = 10.96$ | 0.00563 | 0.02397 | 0.458 | 19 |

### S6 QCM-D of Stce-dSF on ECM models

Quartz crystal microbalance with dissipation (QCM-D) was used to quantify the adsorption and retention of StcE-dSF on immobilized Col-II, Col-I, and FN films. Preparation and characterization of the corresponding ECM precursor films and adsorption of untreated dSF have been reported previously.[2, 8] The QCM-D measurements performed in the present study were therefore designed specifically to determine how StcE treatment alters dSF interactions with the three ECM interfaces. QCM-D measurements were performed using a QCM-I mini (Micro Vacuum, Hungary) and a Q-Sense Explorer microbalance (Biolin Scientific, Sweden) equipped with AT-cut quartz crystals having a fundamental resonance frequency of 5 MHz and a 50 nm gold coating. Measurements were performed at 25 °C. Outlet channels were connected to peristaltic pumps to maintain controlled solution exchange at a flow rate of 200 $\mu L\,min^{-1}$.

#### S6.1 Preparation of ECM-functionalized QCM-D crystals

Gold-coated quartz crystals were rinsed sequentially with 2 wt% SDS, ultrapure water, and ethanol, with the cleaning sequence repeated three times. Crystals were dried under a gentle stream of nitrogen and treated with oxygen plasma (Harrick Plasma, PDC-32G-115 V) for 2 min.

The cleaned crystals were incubated overnight at 4 °C in a 0.088 $mol\,L^{-1}$ ethanolic solution of cysteamine hydrochloride to form amine-terminated self-assembled monolayers (SAMs). The following day, crystals were gently rinsed with ethanol to remove unbound cysteamine and dried under nitrogen. SAM-functionalized crystals were subsequently activated with 25% aqueous glutaraldehyde for 30 min at room temperature, rinsed with ultrapure water, dried under nitrogen, and immediately mounted in the QCM-D fluidic cell.

All ECM precursor films used in the present study were formed at pH 7.4 on cysteamine hydrochloride SAMs. Following establishment of a stable PBS baseline, Col-II, Col-I, or FN at 50 $\mu g\,mL^{-1}$ in PBS at pH 7.4 was introduced into the QCM-D cell. After filling the fluidic chamber, flow was stopped and frequency and dissipation changes were recorded until steady state was reached. This state was defined as the ECM pre-wash condition. Loosely associated protein was subsequently removed by rinsing with PBS at pH 7.4 until a new steady state was reached, defined as the ECM post-wash condition. This preparation follows previously established protocols for Col-II and Col-I and for extended FN immobilized at neutral pH.[2, 8]

#### S6.2 Adsorption of StcE-treated synovial fluid

Following formation and rinsing of the ECM precursor film, non-specific adsorption sites were blocked by exposure to 10 $mg\,mL^{-1}$ bovine serum albumin (BSA) in PBS at pH 7.4 until steady state was reached. The surface was then rinsed with PBS to remove loosely associated BSA.

StcE-treated dSF was subsequently introduced into the QCM-D chamber and allowed to interact with the Col-II, Col-I, or FN precursor film until the frequency and dissipation responses reached steady state. This condition was defined as the StcE-dSF pre-wash state. The experiment was concluded by rinsing the surface with PBS at pH 7.4 until a stable response was obtained. This final state was defined as StcE-dSF post-wash.

The StcE-dSF-associated QCM-D response was calculated relative to the BSA post-wash state so that the contribution of the underlying ECM precursor film and blocking layer was excluded from estimates of StcE-dSF adsorption and retention.

Frequency and dissipation values recorded during the final 2 min of each steady-state step were averaged for subsequent analysis. Each condition consisted of at least 3 independently prepared surfaces.

#### S6.3 QCM-D data analysis and mass modeling

Changes in resonance frequency, $\Delta f_n$, and dissipation, $\Delta D_n$, were recorded simultaneously at the available overtones. QCM-D data were analyzed using pyQCM-BraTaDio.[9] An apparent Sauerbrey areal mass, $m_S$, was estimated from the frequency response according to

$$m_S = -C\frac{\Delta f_n}{n}, \tag{S2}$$

where $C = 17.7\,\mathrm{ng\,cm^{-2}\,Hz^{-1}}$ is the mass-sensitivity constant for a 5 MHz quartz crystal, $\Delta f_n$ is the frequency shift measured at overtone $n$, and $n$ is the overtone number.

Changes in frequency from the available overtones, typically the third through the thirteenth, were analyzed by linear regression using pyQCM-BraTaDio as described previously.[2, 8, 10]

For StcE-treated dSF, adsorption was defined as the apparent Sauerbrey mass measured at the pre-wash steady state relative to the BSA post-wash state, whereas retention was defined as the corresponding mass remaining after the final PBS wash. Because QCM-D detects acoustically coupled solvent in addition to biomolecular material, $m_S$ was treated as an apparent hydrated areal mass. For QCM-D, independent replicates corresponded to separately prepared sensors, whereas MPTM replicates corresponded to independently prepared samples. Measurements acquired at different stages or depths from the same preparation were treated as paired. Data are presented as means with standard error of the mean (SEM), as specified in the figure captions.

#### S6.4 QCM-D statistics

Sauerbrey masses $m_f$ were analyzed before (pre-wash) and after (post-wash) PBS rinsing. Differences between dSF and StcE-dSF on individual ECM substrates were assessed using two-sided Welch's $t$-tests, treating the previously published dSF reference measurements and the StcE-dSF measurements as independent groups. Changes induced by washing were calculated within matched replicates as $\Delta m_f = m_{f,\text{post}} - m_{f,\text{pre}}$. Paired $t$-tests were used for within-experiment wash comparisons. Separate two-factor ANOVAs were performed for pre-wash mass, post-wash mass, and paired wash-induced changes, with ECM identity and fluid treatment as factors. Models employed sum-to-zero contrasts and Type-III partial $F$-tests. Treatment, substrate, and wash-stage comparisons were adjusted using the Holm-Bonferroni procedure within predefined comparison families. Differences in treatment effects between ECM substrates were additionally assessed.

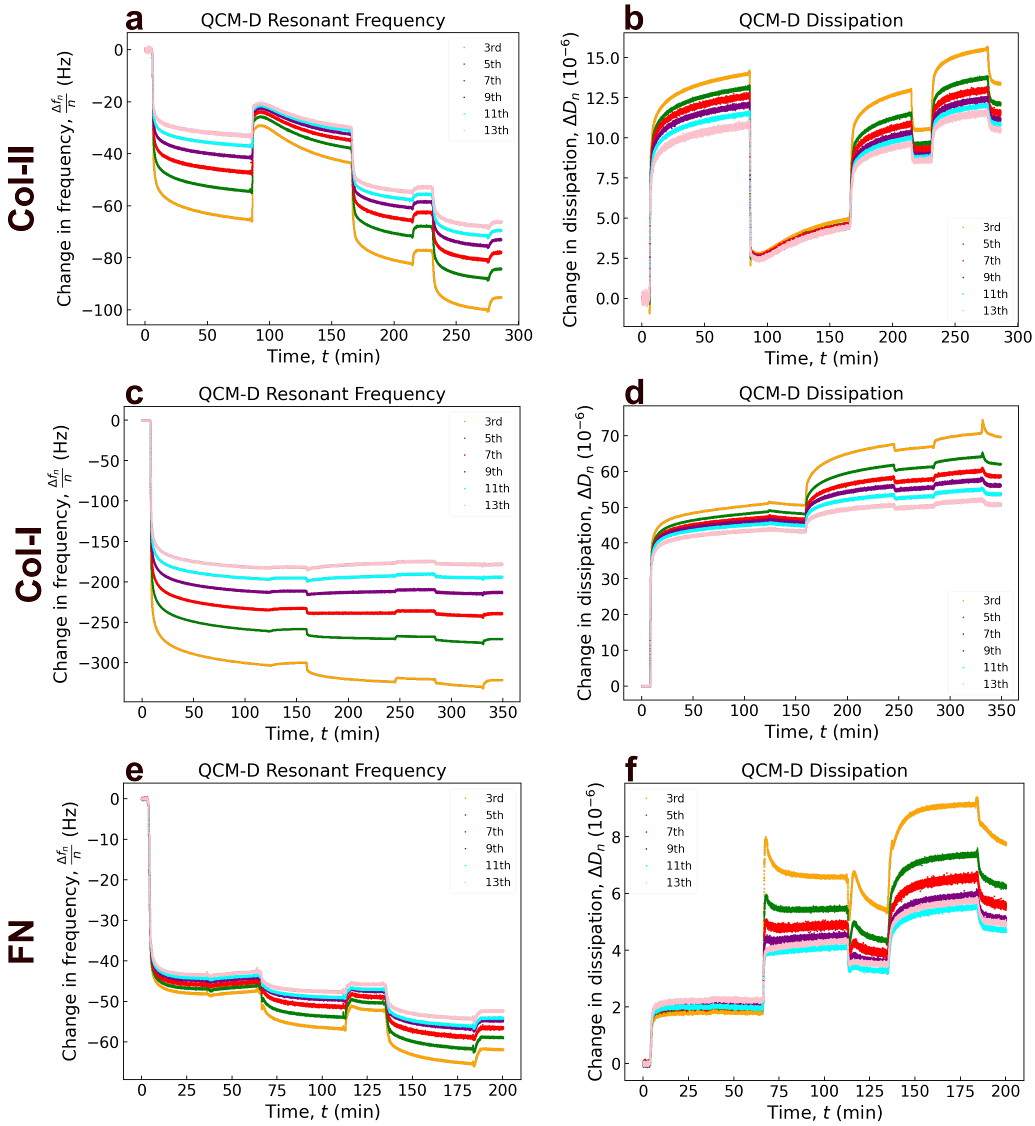


**Figure S7.** Time-resolved overtone-normalized frequency shifts, $\Delta f_n/n$, and dissipation shifts, $\Delta D_n$, during sequential surface functionalization and StcE-dSF exposure on **a,b,** Col-II, **c,d,** Col-I, and **e,f,** FN films. Each experiment consisted of immobilization of the corresponding protein, followed by a PBS rinse, blocking with bovine serum albumin (BSA, 10 mg/mL in PBS), a second PBS rinse, StcE-dSF exposure, and a final PBS rinse to assess the retention of adsorbed components. Traces represent the 3rd, 5th, 7th, 9th, 11th, and 13th overtones.

**a QCM-D apparent mass during exposure and after PBS rinse**

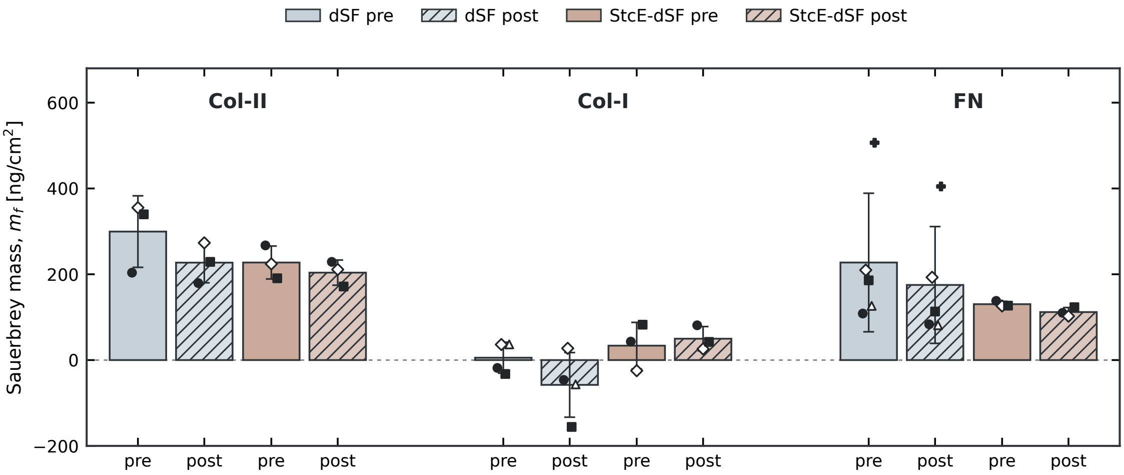


**b Pre-wash planned contrasts**

| Comparison | Δ [95% CI] (ng/cm²) | Holm-adjusted p |
|---|---|---|
| Col-II: dSF vs StcE-dSF | -72 [-247, +103] | 0.7508 ns |
| Col-I: dSF vs StcE-dSF | +28 [-81, +137] | 0.7508 ns |
| FN: dSF vs StcE-dSF | -97 [-297, +103] | 0.7508 ns |
| Col-II vs Col-I (StcE-dSF) | -194 [-305, -82] | **0.0285** * |
| Col-II vs FN (StcE-dSF) | -97 [-189, -5] | 0.0896 ns |
| Col-I vs FN (StcE-dSF) | +97 [-35, +229] | 0.0896 ns |

**c Post-wash planned contrasts**

| Comparison | Δ [95% CI] (ng/cm²) | Holm-adjusted p |
|---|---|---|
| Col-II: dSF vs StcE-dSF | -23 [-119, +73] | 0.7177 ns |
| Col-I: dSF vs StcE-dSF | +108 [-6, +221] | 0.1748 ns |
| FN: dSF vs StcE-dSF | -63 [-232, +105] | 0.7177 ns |
| Col-II vs Col-I (StcE-dSF) | -154 [-219, -89] | **0.0085** ** |
| Col-II vs FN (StcE-dSF) | -92 [-156, -28] | **0.0443** * |
| Col-I vs FN (StcE-dSF) | +62 [+1, +124] | **0.0489** * |

**d Two-way factorial ANOVA: ECM substrate × StcE treatment**

| Stage | Effect | F statistic | p (OLS) | p (HC3) | Partial η² | N |
|---|---|---|---|---|---|---|
| Pre-wash | ECM substrate | F(2,15) = 11.45 | **0.0010** | **<0.0001** | 0.604 | 21 |
| Pre-wash | StcE treatment | F(1,15) = 1.28 | 0.2760 | 0.2289 | 0.079 | 21 |
| Pre-wash | ECM × StcE | F(2,15) = 0.88 | 0.4365 | 0.2810 | 0.105 | 21 |
| Post-wash | ECM substrate | F(2,15) = 12.40 | **0.0007** | **<0.0001** | 0.623 | 21 |
| Post-wash | StcE treatment | F(1,15) = 0.04 | 0.8482 | 0.8219 | 0.003 | 21 |
| Post-wash | ECM × StcE | F(2,15) = 2.12 | 0.1548 | 0.0799 | 0.220 | 21 |

**Figure S8.** QCM-D statistical analysis of untreated and StcE-treated diluted synovial fluid (dSF and StcE-dSF) on extracellular matrix (ECM) substrates. **a,** Sauerbrey mass on Col-II, Col-I, and FN before and after PBS rinsing. Bars represent mean ± SEM; symbols indicate independent measurements, with matching symbols denoting paired pre- and post-wash values. Previously published untreated dSF data are re-plotted and included for comparison.[2, 8] **b,c,** Pre-wash (**b**) and post-wash (**c**) pairwise comparisons between treatments and among StcE-dSF-exposed substrates. Mean differences (Δ) and unadjusted 95% confidence intervals are reported, with Welch's $t$-tests and Holm correction applied within each three-comparison family. **d,** Two-way ANOVA of ECM identity, StcE treatment, and their interaction, reporting $F$-statistics, conventional (OLS) and robust (HC3) $p$-values, partial $\eta^2$, and sample sizes.

### S7 Trackpy Analysis

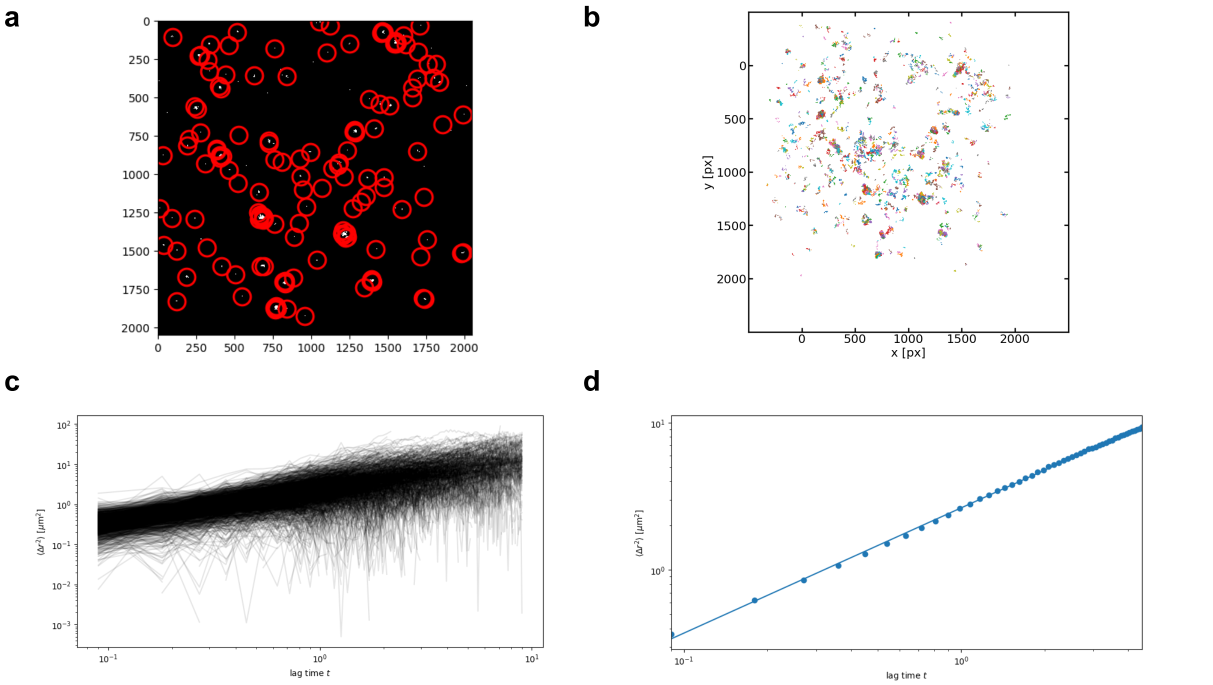


**Figure S9.** Representative workflow of the analysis of tracer-particles for characterization. **a**, Representative frame of tracer-particles in rEqLub at 100 μm from glass. Red circles show detected beads, before filtering. **b**, Trajectories of beads post-filtering and after accounting for drift. **c** All bead displacements shown over their representative lag times, before being averaged together for calculations. **d** Final MSD over lag time plot, with a best fit line plotted, which is used to compute apparent viscosity.

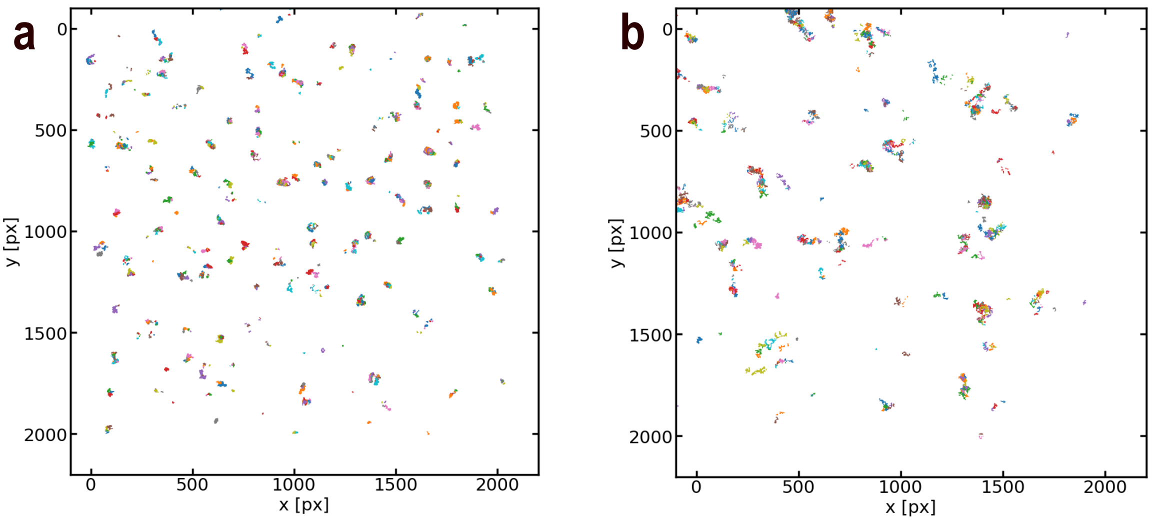


**Figure S10. a,b,** Representative tracer-particle trajectories at 100 μm above glass in dSF (a) and PBS (b), reconstructed using Trackpy and corrected for drift by subtracting the frame-to-frame mean particle displacement. The drift-corrected trajectories were subsequently used to calculate mean-squared displacements (MSDs).

[3] David Nečas and Petr Klapetek. Gwyddion: An open-source software for SPM data analysis, 2 2012.

[4] Von Herrn Heinrich Hertz. Ueber die Berührung fester elastischer Körper. *Journal für die Reine und Angewandte Mathematik*, 1882.

[5] Edward Dintwa, Engelbert Tijskens, and Herman Ramon. On the accuracy of the Hertz model to describe the normal contact of soft elastic spheres. *Granular Matter*, 10(3):209–221, 3 2008.

[6] Krashn Kumar Dwivedi, Piyush Lakhani, Sachin Kumar, and Navin Kumar. Effect of collagen fibre orientation on the Poisson's ratio and stress relaxation of skin: An ex vivo and in vivo study. *Royal Society Open Science*, 9(3), 2022.

[7] Ariell M Smith, Brandon M Pardi, Isaac Sousa, Arvind Gopinath, and Roberto Carlos Andresen Eguiluz. Adenocarcinoma cell mechanobiology is altered by the loss modulus of the surrounding extracellular matrix, 2 2026.

[8] Diego R. Jaramillo Pinto, Nicolas L. Mendoza, Syeda Tajin Ahmed, Yidan Wen, Lenka Vitkova, Spencer M. Witt, Kaleb A. Cutter, Ummay Honey, Matthew J. Paszek, Heidi L. Reesink, Lawrence J. Bonassar, Kevin De France, and Roberto C. Andresen Eguiluz. Functional Differentiation of Type II and Type I Collagen Articular Models in Synovial Fluid Film Formation and Recombinant Equine Lubricin Retention. *Advanced Functional Materials*, 8 2026.

[9] Brandon Pardi, Syeda Tajin Ahmed, Silvia Jonguitud Flores, Warren Flores, Jean-Michel Friedt, Laura L. E. Mears, Bernardo Yáñez Soto, and Roberto C. Andresen Eguiluz. pyQCM-BraTaDio: A tool for visualization, data mining, and modelling of Quartz crystal microbalance with dissipation data. *Journal of Open Source Software*, 9(99):6831, 7 2024.

[10] Sixin Zhai, Diego R. Jaramillo Pinto, Nicolas L. Mendoza, Adekunle Adewole, Benjamin Heufner, Andrea D. Merg, Tomas P. Corrales, Jing Yan, and Roberto C. Andresen Eguiluz. Surface-specific film assembly of a Vibrio cholerae adhesin-derived peptide modulated by environmental salts. *bioRxiv (preprint)*, 6 2026.
